# Sex-specific divergent maturational trajectories in the postnatal rat basolateral amygdala

**DOI:** 10.1101/2021.09.06.458843

**Authors:** Pauline Guily, Olivier Lassalle, Pascale Chavis, Olivier JJ Manzoni

## Abstract

The basolateral amygdala (BLA), the part of the amygdala complex involved in the transduction of perceptual stimuli into emotion, undergoes profound reorganization at adolescence in rodents and humans. How cellular and synaptic plasticity evolve throughout postnatal development in both sexes is only partially understood. We used a cross-sectional approach to compare the morphology, neuronal, and synaptic properties of BLA neurons in rats of both sexes at adolescence and adulthood. While BLA pyramidal neurons from rats of both sexes displayed similar current-voltage relationships, rheobases, and resting potentials during pubescence, differences in these parameters emerged between sexes at adulthood: BLA neurons were more excitable in males than females. During pubescence, BLA neuron excitability was highest in females and unchanged in males; male action potentials were smaller and shorter than females and fast afterhyperpolarizations were larger in males. During post-natal maturation, no difference in spine density was observed between groups or sexes but spine length increased and decreased in females and males, respectively. A reduction in spine head diameter and volume was observed exclusively in females. Basic synaptic properties also displayed sex-specific maturational differences. Stimulus-response relationships and maximal fEPSP amplitudes where higher in male adolescents compared with adults but were similar in females of both ages. Spontaneous excitatory postsynaptic currents mediated by AMPA receptors were smaller in BLA neurons from adolescent female compared with their adult counterparts but were unchanged in males. These differences did not directly convert into changes in overall synaptic strength estimated from the AMPA/NMDA ratio, which was smaller in adolescent females. Finally, the developmental courses of long-term potentiation and depression (LTP, LTD) were sexually dimorphic. LTP was similarly present during the adolescent period in males and females but was not apparent at adulthood in females. In contrast, LTD followed an opposite development: present in adolescent females and expressed in both sexes at adulthood. These data reveal divergent maturational trajectories in the BLA of male and female rats and suggest cellular substrates to the BLA linked sex-specific behaviors at adolescence and adulthood.

## Introduction

The basolateral amygdala (BLA), a limbic area massively connected to cortical and sub cortical structures is at the interface of perception, emotion, and motor behaviors. Its role in the processing of fearful and rewarding stimuli, as well as emotional memory, is now well established (Bocchio et al., 2017; Janak & Tye, 2015; Phelps, 2004). A growing body of evidence indicates sex-differences in these functions (Cahill et al., 2004; Canli et al., 2002; Greiner et al., 2019; Gruene et al., 2015; Lebron-Milad & Milad, 2012). Moreover, the BLA is implicated in the etiology of neuropsychiatric diseases including post-traumatic stress disorder, depression, and anxiety (Daviu et al., 2019; Mahan & Ressler, 2012; Nestler et al., 2002), characterized by their higher prevalence in the female population (Christiansen & Berke, 2020; Kuehner, 2017; McLean et al., 2011). While the study of females’ brains and sex-differences in preclinical research have recently gained momentum (Shansky & Murphy, 2021), sex-differences in the functions of the BLA have mostly been studied in pathological rodent models (Blume et al., 2019; Geary et al., 2021; Guadagno et al., 2020; Przybysz et al., 2021) and the physiological development of the female BLA remains partially understood.

At adolescence, neural networks are rearranged to allow the emergence of novel behaviors and the following transition to adult-specific circuits (Casey et al., 2008; Spear, 2000). Puberty is a particular time window of adolescent neurodevelopment, characterized by the arrival of sexual maturity and accompanied by a profound regulation of sex hormones (Schneider, 2013). Both adolescence and puberty are essential periods of postnatal brain maturation and are characterized by heightened susceptibility to mental disorders (Schneider, 2013). Studies also suggest that sex-differences in depression prevalence emerge, or at least strengthen, during adolescence (Breslau et al., 2017). Additionally, there is consistent evidence indicating that the BLA activity is not mature at adolescence in humans (Scherf et al., 2013), and that puberty influences the maturation of the human amygdala (Bramen et al., 2011; Goddings et al., 2014). However, the magnitude of amygdala reorganization and possible sex differences during this period remain to be fully explored.

In adult rats, sex and estrus differences in excitatory and inhibitory inputs, as well as in the spine density of principal neurons residing in the lateral nucleus (LA) and the basal nucleus (BA) of the BLA have been reported (Blume et al., 2017). Long-term potentiation (LTP) of LABA synapses in both male and female juvenile rats has been described (Bender et al., 2017; Guadagno et al., 2020). LTP and long-term depression (LTD) are both sensitive to sexual hormones in the BLA (Bender et al., 2017; Krężel et al., 2001; Yang et al., 2017), but little is known about sex-specific maturational trajectories of synaptic plasticity between puberty and adulthood. To our knowledge, no study to date has compared neuronal and synaptic properties in the BLA during the pubertal period in both sexes. To address this gap in knowledge, we compared neuronal properties and excitatory transmission in the BLA of pubescent and adult rats of both sexes. Using ex-vivo electrophysiological recordings in rat BLA slices, we report that excitatory neurons undergo complex sex-specific maturational sequence development. Some parameters such as intrinsic properties and action potential properties of neurons exhibit sex-differences only at a specific time of development, whereas other parameters such as basic excitatory transmission and synaptic plasticity display different maturational profiles between male and female rats.

## Material and methods

Further information and requests for resources should be directed to the Lead Contact, Olivier J.J. Manzoni.

### Animals

Animals were treated in compliance with the European Communities Council Directive (86/609/EEC) and the United States NIH Guide for the Care and Use of Laboratory Animals. The French Ethical committee authorized the project APAFIS#26537-2020070812023339. All rats were obtained from Janvier Labs and group-housed (two to four rats per cage) with ad libitum access to food and water. All groups represent data from a minimum of 2 litters. The animal housing room was maintained on a 12 h-day light/dark cycle at constant room temperature (20 ± 1°C) and humidity (60%). Data were collected blind to the estrous cycle unless specified. Female and male rats were classified based on the timing of their pubertal maturation. Male and female rats do not reach puberty at the same time (Schneider, 2013). Thus, the female ages groups were Pubescent 33<P<38 and Adult at 90<P<120. Male groups were: Pubescent 41<P<56 and Adult at 90<P<120 (Supplementary Fig. 1A). All animals were experimentally naïve and used only once.

### Electrophysiology

Male and female rats were anesthetized with isoflurane and killed as previously described (Bara et al., 2018; Borsoi et al., 2019; Scheyer et al., 2019, 2020a, 2020b). The brain was sliced (300 μm) in the coronal plane with a vibratome (Integraslice, Campden Instruments) in a sucrose-based solution at 4°C (in mM as follows: 87 NaCl, 75 sucrose, 25 glucose, 2.5 KCl, 4 MgCl_2_, 0.5 CaCl_2_, 23 NaHCO_3_ and 1.25 NaH_2_PO_4_). Immediately after cutting, slices containing the basolateral amygdala (BLA) were stored for 1h at 32°C in a low-calcium ACSF that contained (in mm) as follows: 130 NaCl, 11 glucose, 2.5 KCl, 2.4 MgCl_2_, 1.2 CaCl_2_, 23 NaHCO_3_, 1.2 NaH_2_PO_4_, and were equilibrated with 95% O2/5% CO_2_ and then held at room temperature until the time of recording.

During the recording, slices were placed in the recording chamber and superfused at 2 ml/min with normal ACSF that contained (in mm): 130 NaCl, 11 glucose, 2.5 KCl, 2.4 MgCl_2_, 2.4 CaCl_2_, 23 NaHCO_3_, 1.2 NaH_2_PO_4_, equilibrated with 95% O_2_/5% CO_2_. All experiments were done at 32°C. The superfusion medium contained picrotoxin (100 μM, Sigma) to block GABA-A receptors, except for low frequency stimulation (LFS) experiments. All drugs were added at the final concentration to the superfusion medium. Field and whole-cell patch-clamp recordings were performed with an Axopatch-200B amplifier, data were low pass filtered at 2kHz, digitized (10 kHz, DigiData 1440A, Axon Instrument), collected using Clampex 10.2 and analyzed using Clampfit 10.2 (all from Molecular Device, Sunnyvale, USA).

Whole-cell patch-clamp of visualized BLA principal neurons and field potential recordings were made as previously described (Bara et al., 2018; Borsoi et al., 2019; Scheyer et al., 2019, 2020a, 2020b) in coronal slices containing the BLA. Neurons were visualized using an upright microscope with infrared illumination. The intracellular solution was based on K^+^ gluconate (in mM: 145 K^+^ gluconate, 3 NaCl, 1 MgCl_2_, 1 EGTA, 0.3 CaCl_2_, 2 Na^2+^ ATP, and 0.3 Na^+^ GTP, 0.2 cAMP, buffered with 10 HEPES). The pH was adjusted to 7.2 and osmolarity to 290 – 300 mOsm. Electrode resistance was 2 – 4 MOhms. Access resistance compensation was not used, and acceptable access resistance was <30 MOhms. The potential reference of the amplifier was adjusted to zero prior to breaking into the cell. Current-voltage (I–V) curves were made by a series of hyperpolarizing to depolarizing 50 pA current steps after breaking into the cell. To determine rheobase and action potential properties a series of depolarizing 10 pA current steps was applied. Membrane resistance was estimated from the I-V curve around resting membrane potential (Bara et al., 2018; Borsoi et al., 2019; Scheyer et al., 2019, 2020a, 2020b). Spontaneous EPSCs (sEPSCs) were recorded at −70mV for at least 10 min as previously described (Kasanetz & Manzoni, 2009; Martin et al., 2016). To determine AMPA/NMDA ratios, the intracellular solution was based on cesium methanesulfonate (Cs^+^Meth, mM): 143 CsMeSO_3_, 5 NaCl, 1 MgCl_2_, 1 EGTA, 0.3 CaCl_2_, 10 HEPES, 2 Na_2_ATP, 0.3 NaGTP and 0.2 cAMP (pH 7.3 and 290-300 mOsm). AMPA/NMDA ratios were calculated by measuring evoked EPSCs (having AMPAR and NMDAR components) at +30 mV. The AMPAR component was isolated by bath application of an NMDAR antagonist (APV; 50 μM, Tocris)(Kasanetz & Manzoni, 2009).

For extracellular field potential recordings, the stimulating electrode was positioned in the lateral amygdala (LA), close to the external capsula (EC), dorsolateral to the recording electrode placed into the adjacent BA (Supplementary Fig. 1B). The glutamatergic nature of the field EPSP (fEPSP) was systematically confirmed at the end of the experiments using the ionotropic glutamate receptor antagonist CNQX (20 μM), which specifically blocked the synaptic component without altering the non-synaptic component. Both fEPSP area and amplitude were analyzed. Stimulation was performed with a glass electrode filled with ACSF and the stimulus intensity was adjusted ~60% of maximal intensity after performing an input–output curve. For LTP experiments, baseline stimulation frequency was set at 0.1 Hz and plasticity was induced by a theta burst stimulation (TBS; 5 trains of 4 pulses at 100 Hz, 200 ms train interval, repeated 4 times at 10 s interval). For LTD experiments, baseline stimulation frequency was set at 0.0666 Hz and plasticity was induced by low frequency stimulation (LFS; 900 pulses at 1 Hz).

### Analysis

Action potential properties were determined from the first spike induced during current clamp experiment. The voltage at which point the dV/dt trace first passed 5 mV/ms was the threshold. fAHP was measured at a local minimum directly following spike’s repolarization within a couple of ms. The mAHP was measured at a local minimum distinct from the fAHP and occurring 15-150 ms after the spike. The frequency and amplitude of sEPSCs were analyzed with Axograph X using a double exponential template: f(t) = exp(-t/rise) + exp(-t/decay) (rise = 0.5 ms and decay = 3 ms). The detection threshold for the events was set at 2.5 times the baseline noise SD. For AMPA/NMDA ratios, NMDAR component was obtained by digital subtraction of the AMPA-EPSC from the dual component EPSC (AMPA/NMDA ratio = peak AMPA-EPSC at +30mV/ peak NMDA-EPSC at +30 mV). For field recording experiments, the magnitude of plasticity was calculated at 30–40 min after induction as percentage of baseline responses. Percent of LTP, referred as LTP (%) = −1x(fEPSP amplitude at baseline (in % of baseline) - fEPSP amplitude at 30-40min (in % of baseline)). Percent of LTD, referred as LTD (%) = fEPSP amplitude at baseline (in % of baseline) - fEPSP amplitude at 30-40min (in % of baseline). The percent of LTD or LTP was calculated and cumulative frequency distribution plots were built for each group (Chaouloff et al., 2007; Martin & Manzoni, 2014).

### Dendritic spines’ reconstruction and quantification

All neurons recorded with Cs^+^Meth solution were also loaded with neurobiotin through patch pipettes. Slices were then fixed overnight in 4% paraformaldehyde, rinsed with PBS and incubated overnight at 4°C with streptavidin-AlexaFluor555 (1:200 in PBS). Slices were mounted for subsequent confocal imaging. All filled neurons corresponded to typical glutamatergic BLA neurons (Washburn & Moises, 1992) (e.g., large cell body, spiny dendrites, pyramidal to stellate shape). Only neurons showing proper filling of the dendritic tree were included in the analysis. Five to eleven dendrites per neuron were analyzed. Dendrites were located approximatively at 70 to 140 μm of the soma. Stack images were acquired using a Zeiss LSM-800 confocal microscope equipped with a 40 oil-immersion objective (NA: 1,4). Frame size was of 512×512 pixels with a x4 zoom. Laser power and photomultiplier gain were adjusted to obtain few pixels with maximum intensity on dendrite shaft and the offset range was tuned to cutoff background noise. Tri-dimensional deconvolution of each stack was performed with Fiji software to compensate the spherical aberration and to correct the z-smear for reliable spine morphology. Point-spread function (PSF) were computed using PSFGenerator plugging (Kirshner et al., 2013) and deconvolution was realized with DeconvolutionLab2 plugin (Sage et al., 2017) using the Richardson-Lucy method. Tridimensional reconstruction and semiautomated analysis were performed with Imaris (Bitplane, Zurich, Switzerland).

### Statistics

Statistical analysis of data was performed with Prism (GraphPad Software) using tests indicated in the main text after outlier subtraction (Grubb’s test, alpha level 0.05). PCA was computed with R (RCore Team (2020). R: A language and environment for statistical computing. R Foundation for Statistical Computing, Vienna, Austria. URL https://www.R-project.org/.). All values are given as mean ±SEM, and statistical significance was set at p<0.05.

## Results

The aim of this study was two-fold. First, we aimed to uncover potential sex-differences at two key moments of development (pubescence and adulthood), and second, we compared maturational trajectories according to sex. Experiments were done in rats of both sexes at pubescent (female 33<P<38; male 41<P<56) and adult stages (at 90<P<120 for both sexes). For the sake of clarity, we chose to present each studied parameter in a way that appropriately highlighted either sex-differences or maturational differences. All complementary statistics can be found in separated Tables.

### Sex-specific differences in intrinsic properties of rat BLA principal neurons emerge in adulthood

Within the BLA, LA and BA principal neurons have distinct neuronal and synaptic properties (Blume et al., 2017). Thus, we purposedly restrained this study to only one of these nuclei and chose the BA. Intrinsic properties are essential features of pyramidal neurons as well as key determinants of synaptic and network properties (Debanne & Poo, 2010). To compare intrinsic firing properties of principal neurons in our experimental groups, we performed patch-clamp recordings of BLA neurons in acute BLA slices obtained from pubescent and adult male or female rats and measured the membrane reaction profiles in response to a series of somatic current steps (Fig. 1, Tab. 1.1-1.2). Recording sites are reported in Supplementary figure 1.

**Figure 1:**
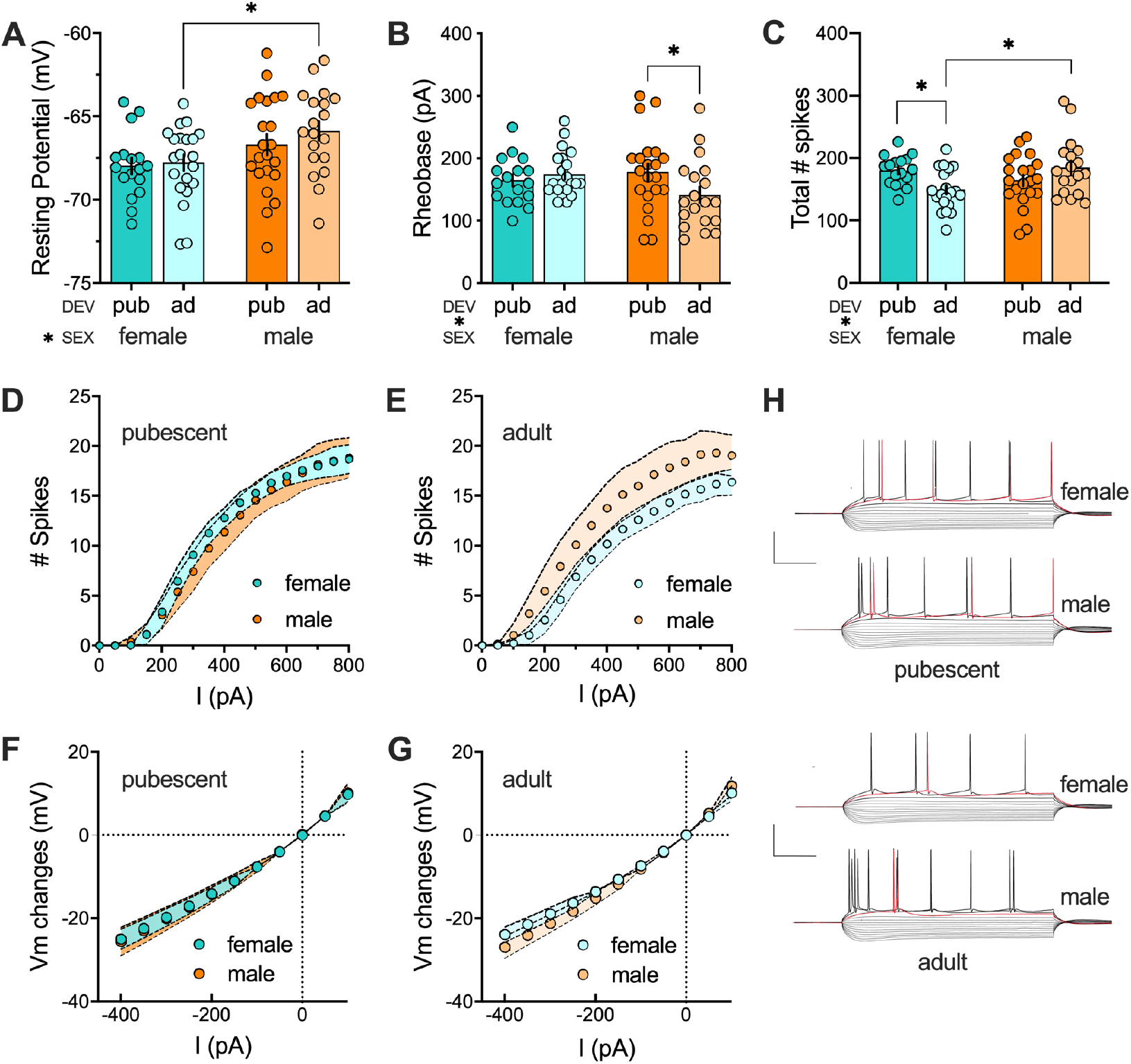
Sex-specific differences in intrinsic properties of rat BLA principal neurons emerge in adulthood. **(A)** Analysis of sex or development main effects and their interaction on the fAHP amplitude revealed that only sex had a significant effect (F (sex 1, 75) = 8.224, P = 0.0054). This effect was driven by significant sex-difference in adult rats, with hyperpolarized neurons in females compared with males (P = 0.0330). **(B)** Progressive current injections in 10pA steps revealed that the minimal current required to trigger an action potential (i.e., rheobase) was influenced by sex and development interaction (F (sex X dev 1, 74) = 4.154, P=0.0451). Post-hoc analysis revealed a higher rheobase in adult males compared with pubescent males (P = 0.0458) while females at different stage of maturation did not differ (P = 0.8036). **(C)** Analysis of the total number of evoked action potentials in response to increasing depolarizing current revealed a significant sex X development interaction F (sex X dev 1, 73) = 9.085, P=0.0035). Adult females spiking less than both pubescent females (P = 0.0276) and adult males (P = 0.0078). **(D-E)** Step by step analysis of the number of evoked action potentials in response to increasing depolarizing current was similar in pubescent rats of both sexes while neurons measured in adult males spiked more than of those of adult females. **(F-G)** Current–voltage (I-V) curves from BLA pyramidal neurons were similar in pubescent and adult rats of both sexes. **(H)** Typical membrane response to current steps of 50 pA from −400 pA to 250 pA in neurons of pubescent and adult rats of both sexes. Data are shown as mean ± SEM (A-C) or mean ± 95% of CI (D-G). Pubescent: female, n = 17/10; male, n = 21-22/10-11. Adult: female, n = 21/12; male, n = 18-19/9-10. n = cells/rats. Data analyzed via 2-way ANOVA followed by Šídák multiple comparison test (A-C). *P<0.05. Color code: females, dark (pubescent) or light (adult) blue; males, dark (pubescent) or light (adult) orange.

At pubescence, all recorded BLA pyramidal neurons showed similar and superimposable I–V plots independent of sex (Fig. 1F). Similarly, the resting membrane potential (Fig. 1A) and the rheobase (Fig. 1B) were similar within and between sex groups. In females but not males, the number of depolarization-driven spikes was larger at adolescence than adulthood (Fig. 1C-D and Tables 1.1-1.2).

At adulthood however, differences emerged between sexes: there was significant, albeit small, shift between I–V plots of males and females (Fig. 1E). Additionally, the resting membrane potential (Fig. 1A) of BLA pyramidal neurons significantly differed between sexes with male neurons being more depolarized than females. Furthermore, as estimated from the number of depolarization-driven spikes, adult males showed more action potentials in response to somatic current steps, compared with females (Fig. 1C and Tables 1.1-1.2).

Further, Principal Component Analysis (PCA, Fig. 2) with resting membrane potentials, rheobase and total number of spikes as quantitative variables, indicated that sex is the principal contributor to the data set variance when all groups are considered. PCA also showed that the sex difference was the most pronounced during adulthood (Fig. 2C).

**Figure 2:**
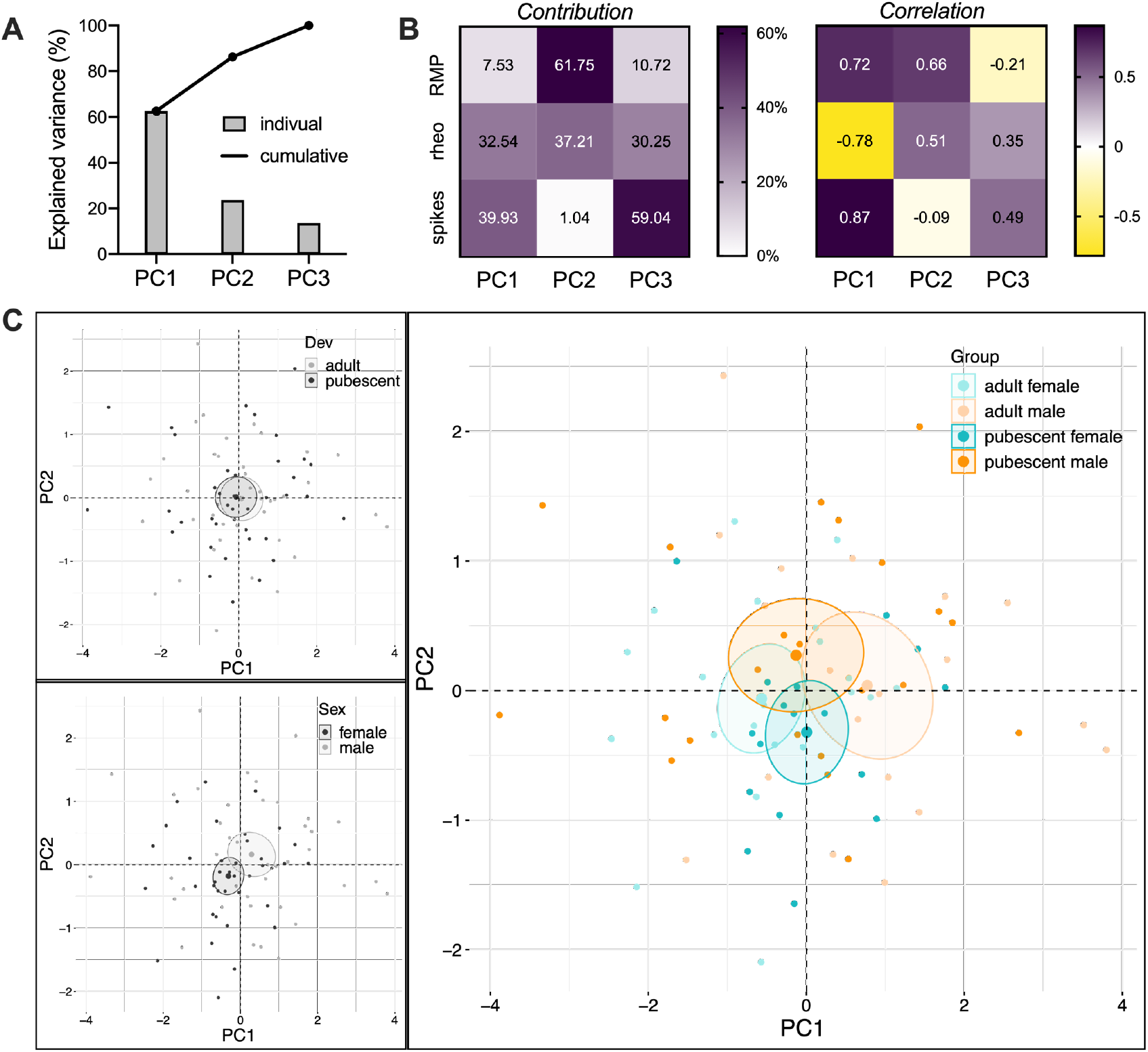
PCA of intrinsic properties show that sex is the principal contributor to the data set variance. Data were analyzed via PCA with resting membrane potential (RMP), rheobase (rheo) and total number of spikes (spikes) as quantitative active variables and individual cells as individuals. Supplementary qualitive variables were development stage (dev, 2 modalities), sex (2 modalities) and group (4 modalities). **(A)** Plotting the percentage of explained variance by each PC (histogram) reveals that most of the data set’s variance is explained by PC1 (62.6 %), followed by PC2 (23.7 %) and PC3 (13.7%). The cumulative percentage of explained is represented by black dots. **(B)** Heatmaps representing the contribution (left) and correlation (right) of each variable with PCs 1, 2 and 3. (Left) Variables with the strongest contribution for a PC were used to build the PC (**C**) and individuals’ coordinate on this axis provides information on the value taken by the individual for this variable. (Right) A positive correlation of a variable with a PC means that individuals that have high or low coordinates on the PC tend to have respectively high or low values for the variable. Conversely, a negative correlation means that individuals that have high coordinates on the PC tend to have low values for the variable or vice et versa. **(C)** PCA graph of individuals was built with PC1 and PC2 which together explained more than 80% of the variance (see A). Small dots represent individuals colored according to their belonging to one the following qualitative supplementary variables: dev (top left), sex (bottom left), group (right). Bigger dots represent the barycenter of individuals (ie. mean) for each category, surrounded by its 95% confidence ellipses (CE). PCA show that pubescents and adults CEs largely overlap (top left). In contrast, males and females CEs overlap only very partially (bottom left), in support of a large contribution of sex in the overall variance of intrinsic properties. Group analysis revealed that this effect is driven by a sex difference in adults (right). Cells obtained from adult males tend to spike more and have a lower rheobase (higher PC1 coordinates) than adult females (lower PC1 coordinates).

Together these data show that sex-specific differences in intrinsic properties emerge at adulthood in BLA pyramidal neurons and that male BLA neurons display a higher excitability than female neurons at the same age.

### Action potentials in BLA principal neurons diverge at pubescence

Changes in spike trains and excitability could partly be explained by changes in ion channel conductance which generate the different phases of action potentials (Hunsberger & Mynlieff, 2020; Matos et al., 2020; Y. Zhang et al., 2020). To gain insight into this matter, we next explored waveform properties of individual action potentials (Fig. 3, Tab. 2.1-2.2). While action potentials’ threshold (Fig.3B), amplitude (Fig.3C), half-width (Fig. 3E), duration (Fig. 3F) and depolarization duration (Fig. 3G) did not vary between our four groups, sex-differences in other action potential properties were found in pubescent rats. Thus, at pubescence, action potentials’ overshoot (Fig. 3D) and repolarization duration (Fig. 3H) were larger in females compared with males, supporting the existence of sex-differences in the repolarization phase of pubescent rat. In females, there was a sex-specific maturation of repolarization duration that was shorter at adulthood (Fig. 3H).

**Figure 3:**
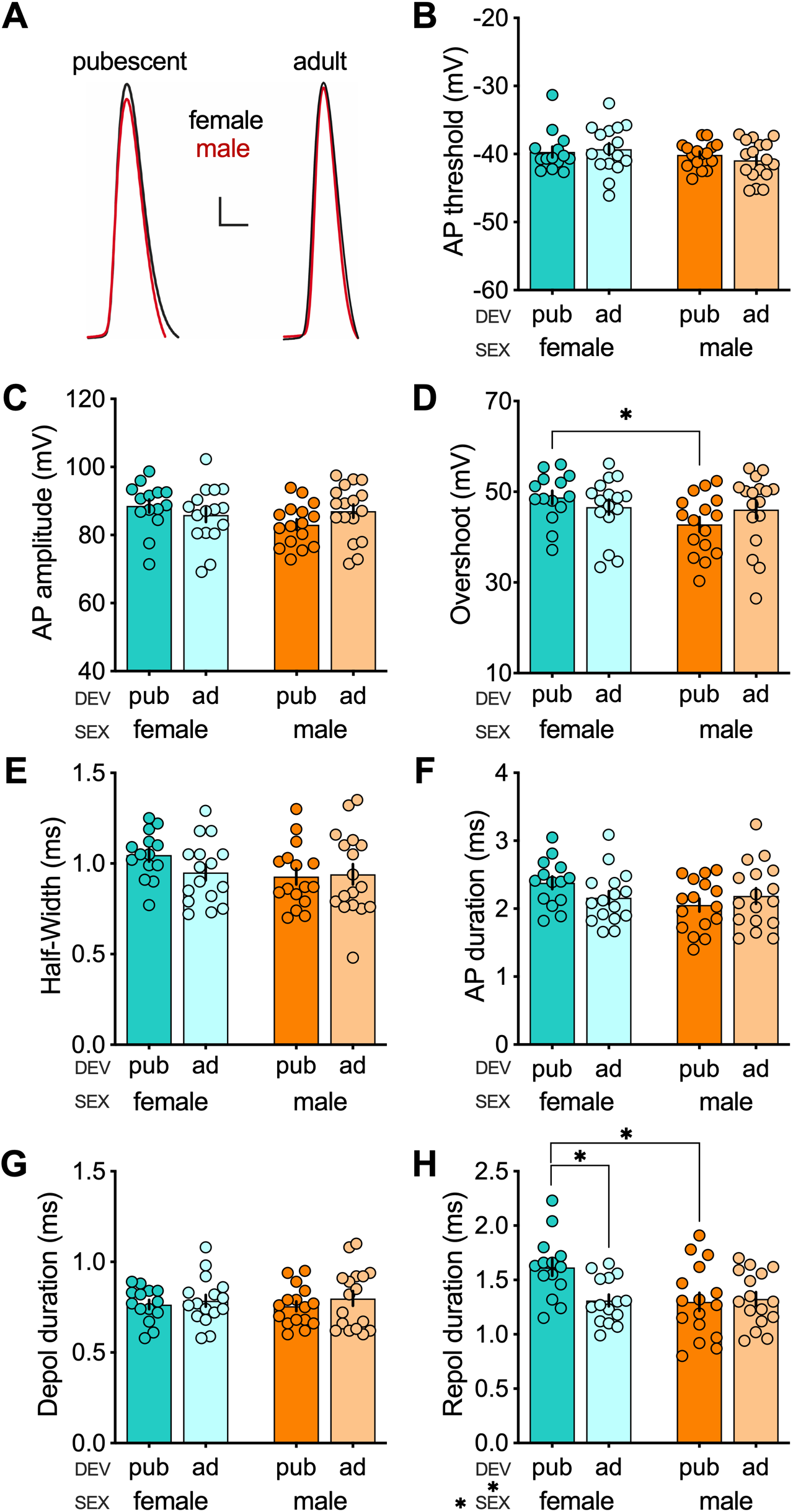
Action potential properties of principal neurons in the rat BLA. **(A)** Representative traces of the first spike elicited by a 10 pA current step injection in neurons of pubescent (right) and adult (left) rats. Male (red) and female (black) traces are superimposed for each developmental stage. Scale bar :10 mV, 1ms. **(B)** Action potential threshold did not vary between groups. **(C-D)** Despite no difference in action potential amplitude (C), the overshoot was lower in pubescent males compared with females (D; P = 0.0392). This sex-difference was not found in adults (P = 0.9685). However, sex (F (sex 1, 59) = 3.573 P=0.0637) and development (F (dev 1, 59) = 0.1067 P=0.7451) main effects, or sex X development interaction (F (sex X dev 1, 59) = 2.495 P=0.1195), did not reach significance. **(E-F)** Half-width (E) and action potential duration (F) did not vary between groups. **(G-H)** Rise time was similar between each group (G). Conversely, the analysis of repolarization duration revealed a significant sex X development interaction (F (sex X development 1, 56) = 6.918 P=0.0110). Repolarization duration was shorter in pubescent males compared with females (P = 0.0025) whereas no difference was found between adult of both sexes (P = 0.9527). Pubescent females also presented a longer repolarization duration compared with adult females (P = 0.0152), whereas no difference was found between males of both ages (P = 0.6103). All data are shown as mean ± SEM. Pubescent: female, n = 14/9; male, n = 16/10. Adult: female, n = 16/7; male = 17/10. n=cells/rats. Data analyzed via 2-way ANOVA followed by Šídák multiple comparison test. *P<0.05. Color code: females, dark (pubescent) or light (adult) blue; males, dark (pubescent) or light (adult) orange.

At adulthood, action potentials of BLA principal neurons displayed similar amplitude (Fig. 3C), overshoot (Fig. 3D), half-width (Fig. 3E) and duration (Fig. 3F).

Rather counter intuitively, these data suggest that, in contrast to sex-differences in other intrinsic properties, sex-differences in action potential properties are only found at pubescence and disappear in adulthood in BLA principal neurons.

In continuity with the intrinsic properties (see Fig. 1 & 2), PCA indicated that again, sex is the principal contributor to the variance of this data set (Fig. 4).

**Figure 4:**
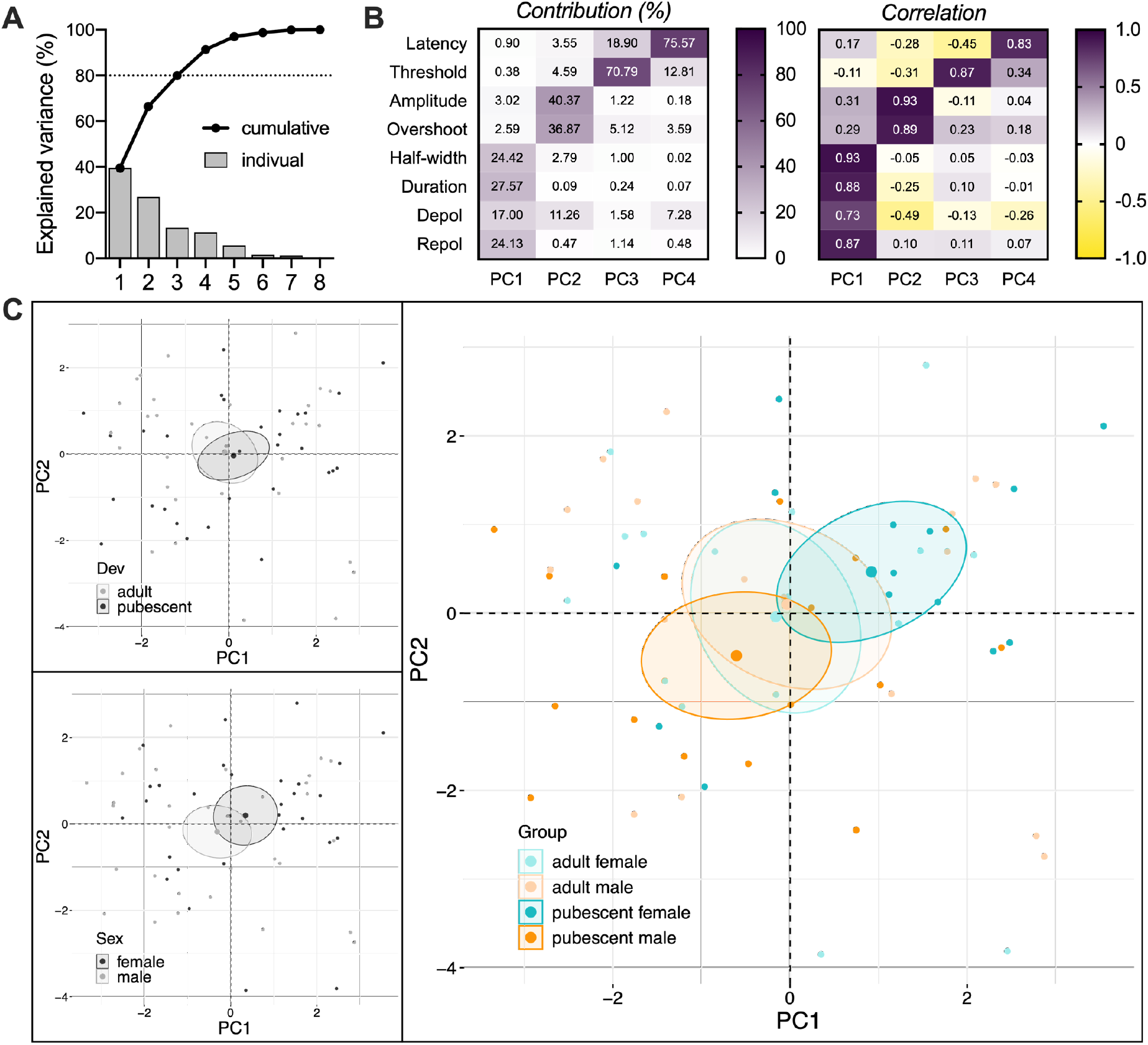
PCA of action potentials’ properties suggest that sex is a major contributor to the data set variance. Data were analyzed via PCA with action potential latency, threshold, amplitude, overshoot, half-width, duration, depolarization duration (depol) and repolarization duration (repol) as quantitative active variables and individual cells as individuals. Supplementary qualitive variables were development stage (dev, 2 modalities), sex (2 modalities) and group (4 modalities). **(A)** Plotting the cumulative relative contribution of PCs against the variance reveals that most of the data set’s variance is explained by PC1 (39,5 %), followed by PC2 (27,0 %), PC3 (13,5 %) and PC4 (11,4 %). **(B)** Heatmaps representing the contribution (left) and correlation (right) of each variable with PCs 1, 2, 3 and 4. (Left) Variables with the strongest contribution for a PC were used to build the PC (**C**) and individuals’ coordinate on this axis provides information on the value taken by the individual for this variable. (Right) A positive correlation of a variable with a PC means that individuals that have high or low coordinates on the PC tend to have respectively high or low values for the variable. Conversely, a negative correlation means that individuals that have high coordinates on the PC tend to have low values for the variable or vice et versa. **(C)** PCA graph of individuals. Small dots represent individuals colored according to their belonging to one the following qualitative supplementary variables: dev (top left), sex (bottom left), group (right). Bigger dots represent the barycenter of individuals (i.e., mean) for each category, surrounded by its 95% confidence ellipses (CE). PCA show that pubescents and adults CEs largely overlap (top left). Males and females CEs overlap only partially (bottom left), in support of a substantial contribution of sex in the overall variance of action potentials’ properties. Group analysis revealed the importance of a sex difference in pubescent in this effect (right). Cells obtained from pubescent females have longer and larger PAs (higher PC1 and PC2 coordinates, respectively) than pubescent males.

The prolonged activation of calcium-dependent potassium channels (KCa) during the repolarization phase of action potentials leads to an afterhyperpolarization (AHP) potential below the action potential threshold. The medium AHP (mAHP, Fig. 5A), is tightly linked to neuronal excitability and sensitive to stress exposure in the BLA (Faber et al., 2001; Hetzel & Rosenkranz, 2014; Rau et al., 2015). Less is known about the fast AHP (fAHP, Fig. 5A), however, it undergoes a profound maturation within the first post-natal month in the BLA (Ehrlich et al., 2012). The amplitudes of the fAHP and mAHP were measured in the first action potential recorded at the rheobase (Fig. 5, Tab. 3.1-3.2).

**Figure 5:**
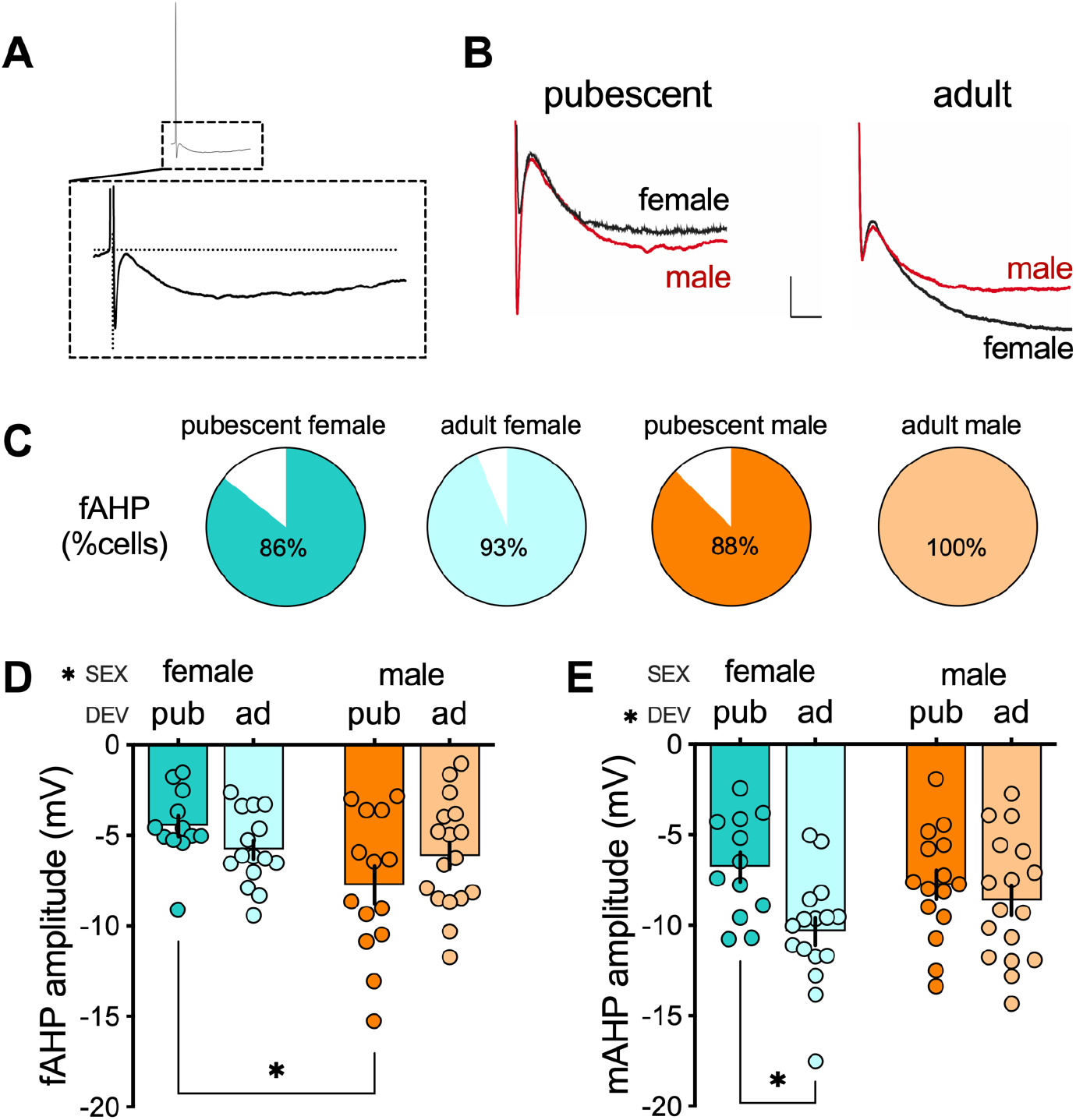
AHP profile of principal neurons in the rat BLA of pubescent and adult rat of both sexes. **(A)** Bottom: zoomed representation of the AHP of the action potential on the top. The fAHP is the local minimum directly following spike’s repolarization within a couple of ms. The mAHP is a local minimum distinct from the fAHP and occurring 15-150 ms after the spike. **(B)** Representative traces of the first spike’s AHP elicited by a 10 pA current step injection in neurons of pubescent (left) and adult (right) rats. Male (red) and female (black) traces are superimposed for each development stage group. Scale bar: 2 mV, 20ms. **(C)** Proportion of cells with a first spike expressing a fAHP in pubescent (female, n = 12/14 cells, 86%; male, n = 14/16 cells, 88%,) and adult (female, n = 15/16 cells, 93%; male, n = 17/17 cells, 100%) rats. **(D)** Analysis of sex or development main effects and their interaction on the fAHP amplitude revealed that only sex had a significant effect (F (sex 1, 54) = 5.708, P = 0.0204). This effect was driven by significantly larger fAHPs in pubescent males compared with females (P = 0.0106). Pubescent: female, n = 12/9; male, n = 14/10. Adult: female, n = 15/8; male, n = 17/10. n = cells/rats. **(E)** The mAHP amplitude was influenced by development (F (development 1, 56) = 7.364 P = 0.0088). This main effect was driven by a significantly larger mAHP in adult females compared with pubescent females (P = 0.0084). Pubescent: female, n = 12/8; male, n = 15/9. Adult: female, n = 16/8; male, n = 17/10. n = cells/rats. All data are shown as mean ± SEM. Data analyzed via 2-way ANOVA followed by Šídák multiple comparison tests. *P<0.05. Color code: females, dark (pubescent) or light (adult) blue; males, dark (pubescent) or light (adult) orange.

In every group, a large majority of neurons displayed a fAHP (Fig. 5C). There was a significant effect of sex (P=0.0204) and only a trend toward a significant effect of interaction between age and sex factors (P=0.0590). These effects were driven by a significantly greater fAHP amplitude pubescent males compared with females. Note that the latter sex-difference was no longer present in adults (Fig 5D). Further, there was a maturation of the mAHP in females, illustrated by a larger mAHP in adult compared with their pubescent counterparts (Fig. 5E).

### Synaptic properties in the BLA of pubescent and adult rats

Synaptic properties were compared and input–output curves constructed from the fEPSP amplitude vs increasing stimulation intensities at the BLA from brain slices of our 4 groups (Fig. 6, Tab. 4.1-4.2). Significant differences in stimulus-response relationship and maximal fEPSP amplitude were found between pubescent and adult males (Fig. 6B-C). In females, the stimulus-response relationship did significantly vary between pubescents and adults (Fig. 6A), however the maximal fEPSP amplitudes did not (Fig. 6C), despite a propensity towards larger fEPSP in pubescent females.

**Figure 6:**
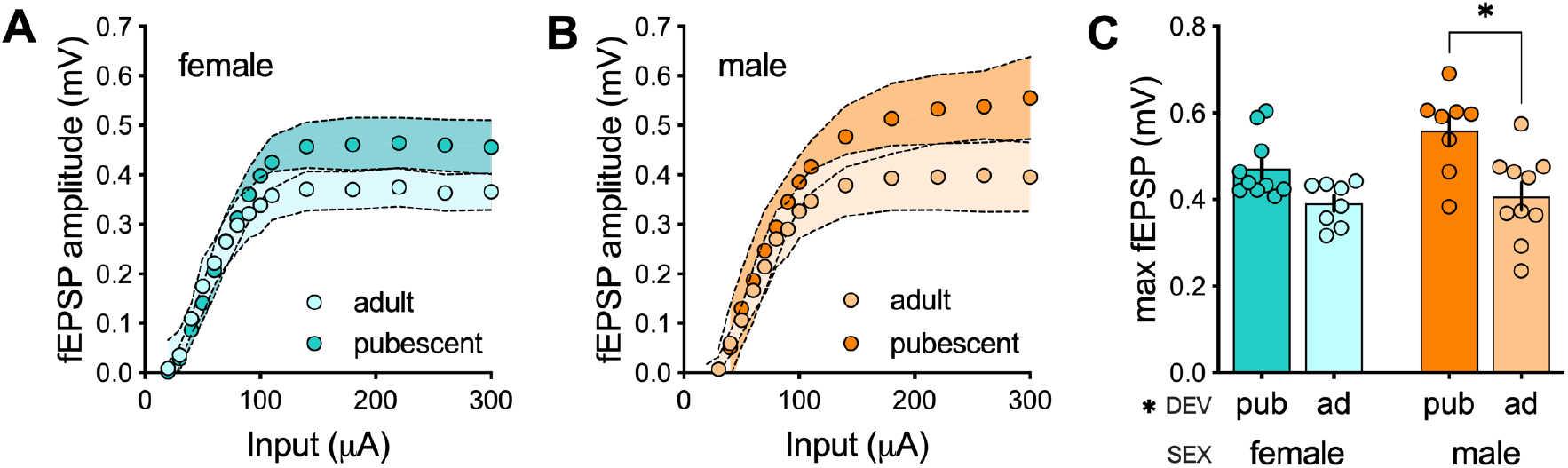
fEPSPs input-output profile. **(A-B)** Averaged fEPSP amplitude as a function of the stimulus intensity. There is a maturation of the input-output profile between pubescence and adulthood in both females (A) and males (B). **(C)** Development influenced the maximum amplitude of fEPSPs, pubescent rats having larger fEPSPs than adults (F (development 1, 32) = 17.59 P = 0.0002). This effect was driven by larger maximum fEPSPs in pubescents males compared with adults (P = 0.0010) while comparison between females of both developmental stages did not reach significance (P = 0.0943). n = individual rats, one slice per rat. Females: pubescent, n = 10; adult, n = 8. Males: pubescent, n = 10; adult, n = 8. n=cells/rats. Data are shown as mean ±CI (A-B) or ± SEM (C). Data analyzed via 2-way ANOVA followed by Šídák multiple comparisons test (C). *P<0.05. Color code: females, dark (pubescent) or light (adult) blue; males, dark (pubescent) or light (adult) orange.

At excitatory synapses, the relative levels of AMPAR and NMDAR is a measure of synaptic integration and plasticity (Malinow & Malenka, 2002). We compared the ratio between AMPAR- and NMDAR-evoked EPSCs (AMPA/NMDA ratio) in adolescent and adult BLA neurons from rats of both sexes (Fig. 7, Tab. 5,1-5,2). This parameter significantly varied between males and females at pubescence (Fig. 7A), males having a greater AMPA/NMDA ratio. In BLA slices of pubescent males, the bimodal distribution of the data clearly shed light on two subgroups of neurons: one group displayed AMPA/NMDA ratio like females and the other exhibited AMPA/NMDA ratios higher than females (Fig 7C). In adult rats, the AMPA/NMDA ratio was similar across sexes (Fig. 7B-D).

**Figure 7:**
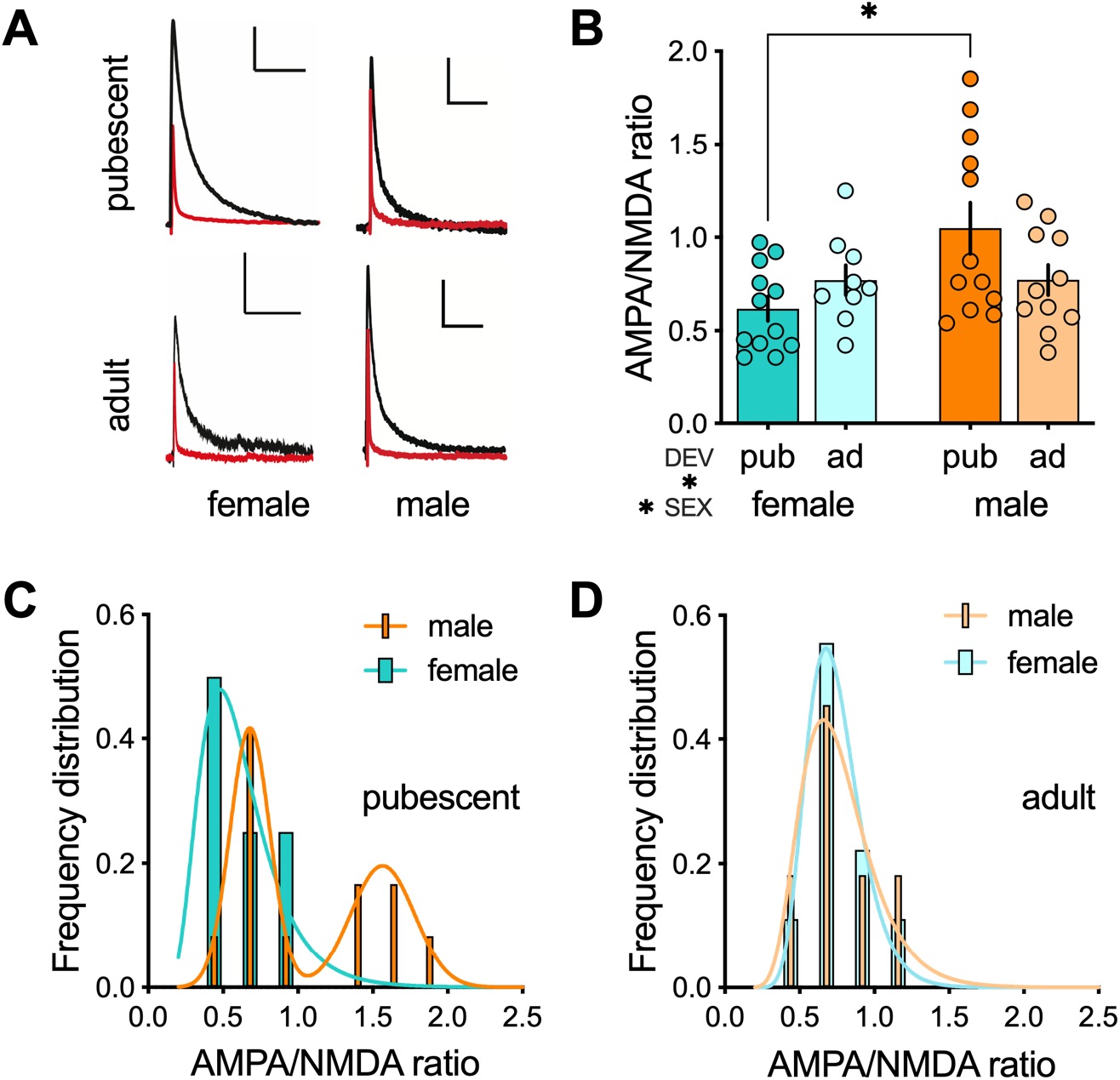
Sex-differences in AMPA/NMDA ratios disappear in adulthood. **(A)** Representative waveforms of AMPA EPSC evoked at +30mV in the presence of 50 μM APV (red) and NMDA EPSC obtained by digital subtraction of the AMPA component from the dual component EPSC evoked at +30mV (black). Scale bar: 500ms, 50 pA. **(B)** The analysis of AMPA/NMDA ratio revealed a significant sex X development interaction (F (sex X development 1, 40) = 4.739 P=0.0355). The AMPA/NMDA ratio of principal neurons was significantly higher in BLA slices obtained from pubescent males, as compared with those obtained from adult pubescent females (P = 0.0048). Conversely, no sex-differences were found in adults (P > 0.999). **(C)** Frequency distribution of AMPA/NMDA ratio reveals a subset of neurons from pubescent males with AMPA/NMDA ratio comparable to pubescent females, and another population of neurons with a greater AMPA/NMDA ratio. AMPA/NMDA ratio data from pubescent females were best fit by a single Gaussian (blue line), whereas data from pubescent males were best fit by the sum of two Gaussians (orange line), indicative of two populations. **(D)** AMPA/NMDA ratio are distributed similarly between female and male adult rats. AMPA/NMDA ratio data of both groups best fit single gaussians. All data are shown as mean ± SEM. Pubescent: female, n = 12/6; male, n = 12/9. Adult, female, n = 9/6; male, n = 11/7. n=cells/rats. Data were analyzed via 2-way ANOVA followed by Šídák multiple comparisons test (B). *P<0.05. Color code: females, dark (pubescent) or light (adult) blue; males, dark (pubescent) or light (adult) orange.

To extend our portrait of excitatory BLA synapses, AMPAR spontaneous EPSCs (sEPSCs) were isolated in BLA neurons voltage-clamped at −70 mV (Fig. 8, Tab. 6,1-6,2). Whereas the frequency of AMPAR sEPSCs remained invariable in females (Fig. 8B), the distribution of sEPSC amplitudes was shifted toward lower values in adolescent female rats (Fig. 8C-D). In contrast, both AMPA sEPSCs amplitude and frequency remained invariable in male littermates (Fig. 8E-H).

**Figure 8:**
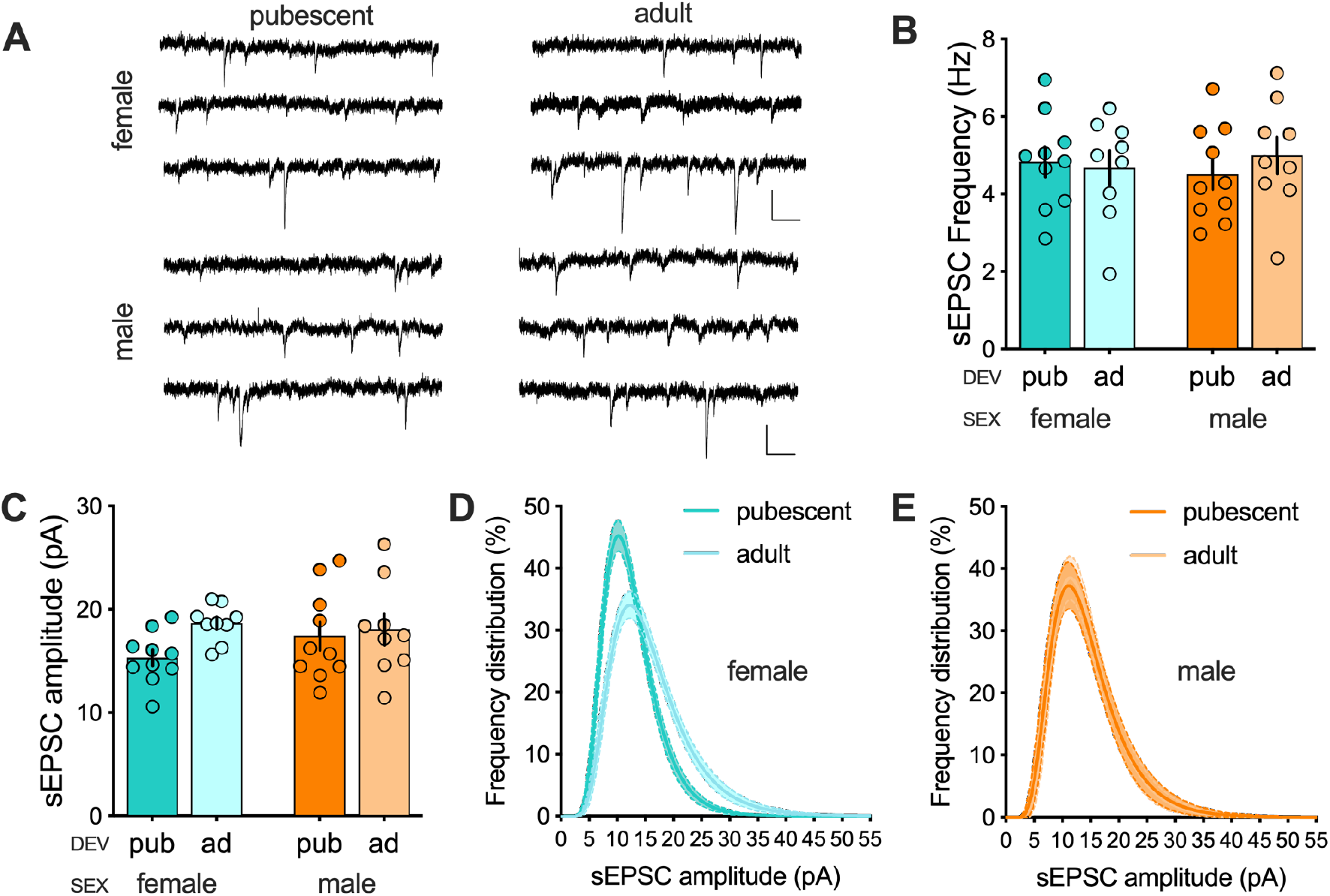
Amplitude distribution of sEPSCs is skewed towards larger amplitudes in adult females but not males BLA pyramidal neurons. **(A)** Representative traces of sEPSCs recorded at −70 mV in BLA pyramidal neurons. Scale bar: **(B)** sEPSCs’ mean frequency was similar for every group. There was no significant effect of development, sex, or their interaction, neither of any multiple comparison tests. **(C)** sEPSCs’ mean amplitude was similar for every group. There was no significant effect of development, sex, or their interaction, neither of any multiple comparison tests. **(D)** Log-normal curve fit (± CI) of frequency distribution of individual sEPSCs’ amplitude reveals a higher proportion of large size events in adult compared to pubescent female rats. **(E)** Individual sEPSCs’ amplitudes are distributed similarly in pubescent and adult males. Lognormal curve fit showing distributions’ similar skewness between pubescent and adult males. All data are shown as mean ± SEM. Females: pubescent, n = 10/8; adult, n = 9/7. Males: pubescent, n = 10/7; adult, 9/7. N = cells/rats. Data were analyzed via 2-way ANOVA followed by Šídák multiple comparisons test (B-C), or 2-way ANOVA of repeated measures followed by Tukey multiple comparisons test for each sex (D-E). *P<0.05. Color code: females, dark (pubescent) or light (adult) blue; males, dark (pubescent) or light (adult) orange.

### Sex specific maturation of BLA neurons dendritic spines

Dendrites of BLA principal neurons are characterized by the presence of numerous spines which are the main contact of glutamatergic inputs (Brinley-Reed et al., 1995; Farb et al., 1992; Radley et al., 2007; Washburn & Moises, 1992). These spines are subject to intense maturational processes in the postnatal brain (Moyer & Zuo, 2018) and the BLA (Bosch & Ehrlich, 2015). Moreover, sex-differences in spine density have also been reported in the adult BLA (Blume et al., 2017). Thus, to elucidate the sex-specific maturation of excitatory synapses in BLA principal neurons, we performed an ex vivo three-dimensional reconstruction of neurobiotin-filled BLA neurons (Fig. 9, Tab. 7.1-7.4).

**Figure 9:**
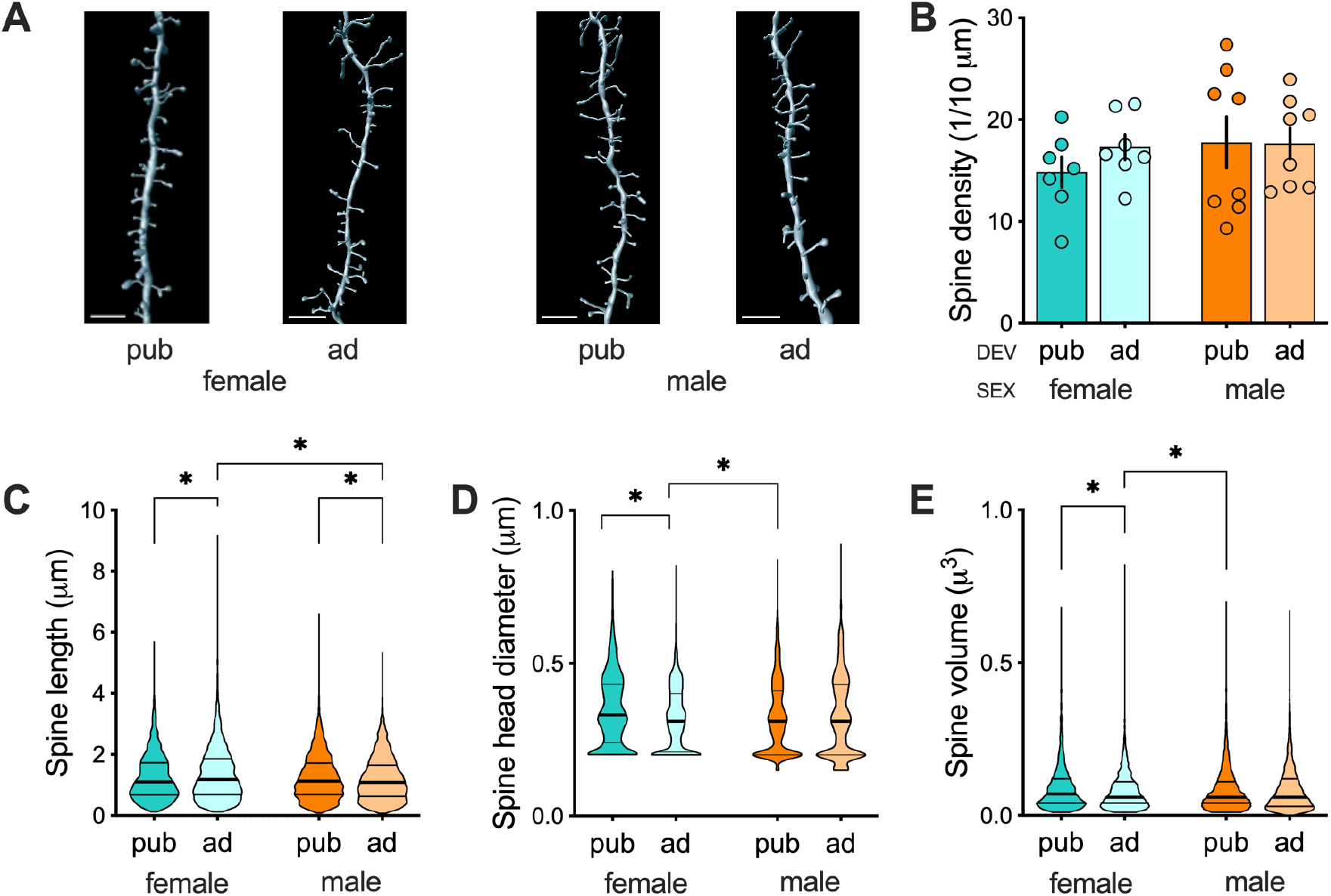
Sex-specific dendritic spine maturation of BLA principal neurons. **(A)** Representative 3D rendering of dendritic spines from neurobiotin filled principal neurons. Scale bar: 4μm. **(B)** Dendritic spine density was similar in all the groups. There was no significant effect of development, sex or their interaction, neither of any multiple comparison test. Females: pubescent, n = 7/7; adult, n = 7/4. Males: pubescent, n = 8/6; adult, n = 8/5. n = cells/rats. **(C)** For neurons obtained from females, dendritic spines were shorter in pubescent compared with adult rats (P=0.0061), whereas the opposite was found in males, with longer spines in pubescent compared with adult rats (P=0.0018). Adult females also had longer spines compared with adult males (P<0.0001). Females: pubescent, n = 2450; adult n = 3908. Males: pubescent, n = 4658; adult, n = 4639. n = spines. **(D)** In females, spines’ heads diameters were larger in pubescent compared with adult rats (P<0.0001), whereas in males no significant development difference in spines’ heads diameters was found (P>0.9999). Pubescent females also had spines with larger heads compared with pubescent males (P<0.0001). Females: pubescent, n = 2450; adult n = 3784. Males: pubescent, n = 4640; adult, n = 4585. n = spines. **(E)** Spine volumes were bigger in pubescent females compared with both adult females and pubescent males. Females: pubescent, n = 2430; adult n = 3769. Males: pubescent, n = 4586; adult, n = 4540. n = spines. Data are shown as mean ± SEM (B) or violin plot with median (center), interquartile ranges (bounds), maxima and minima (C-E). Data were analyzed via 2-way ANOVA followed by Šídák multiple comparisons test (B) or Kruskal-Wallis test followed by Dunn’s multiple comparisons test (C-E). Color code: females, dark (pubescent) or light (adult) blue; males, dark (pubescent) or light (adult) orange.

Quantitative analysis of dendritic spine density and architecture revealed no change in the density of spines in our different groups (Fig. 9A-B; all classes of spines were analyzed). Analyzing the distribution of spine length, spine head diameter, and spine volume, we found significant differences between adult and pubescent spines of sex-matched animals (Fig. 9C-E). Analyzing the total population of spines revealed a more pronounced maturation in females than males. Specifically, there was an augmentation in spine length (Fig. 9C), paralleled by a reduction in spine head diameter (Fig. 9D) and spine volume (Fig. 9E) in adult females compared with pubescents females. In males, differences were restricted to a reduction in the spine length of adults compared with pubescent rats (Fig. 9C). Sex differences manifested at pubescence: Spine head diameters and volumes were bigger in pubescent females than males (Fig. 9D and E, respectively).

### Theta-burst long-term potentiation follows a sex-specific maturational sequence in the rat BLA

Long-term potentiation (LTP) of excitatory synapses has been extensively studied in the BLA due to its role in fear learning and memory (Pape & Pare, 2010). While most studies measured LTP at sensory inputs onto the LA, LTP at LA-BA synapses has also been reported in male and female rats at young ages (Bender et al., 2017; Guadagno et al., 2020). We elected to investigate LA-BA LTP in the BLA of male and female rats at our two age periods to test the hypothesis of a sexual dimorphism of BLA plasticity. We chose a mild induction protocol (theta burst stimulation, TBS) that allows detecting of altered LTP in rodent models of neuropsychiatric disorders (Iafrati et al., 2016; Labouesse et al., 2017; Manduca et al., 2017; Thomazeau et al., 2014). The LTP-induction protocol was applied to acute BLA slices obtained from our various rat groups during simultaneous recording of extracellular field EPSPs (Fig. 10, Tab. 8).

**Figure 10:**
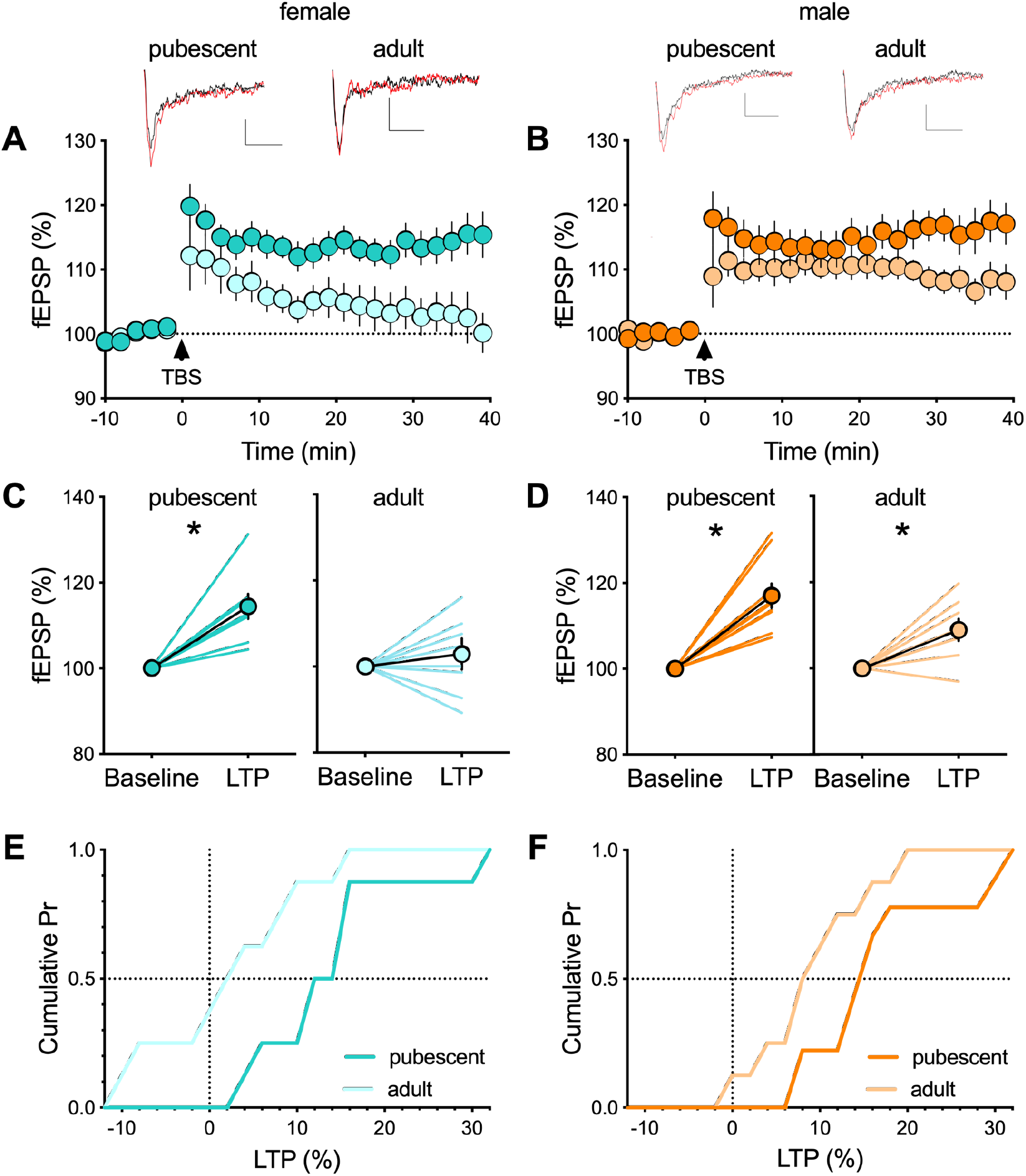
Age- and sex-dependent ablation of long-term potentiation (LTP) in the rat BLA. **(A-B)** Average time-courses of mean fEPSPs amplitude showing that theta-burst stimulation (TBS, indicated arrow) induced LTP at BLA synapses in pubescent females but not in adult females (A). In contrast, LTP was present in BLA slices obtained from both pubescent and adult male rats (B). Above: example traces, baseline (black) and 40 min post-stimulation (red). Scale bar: 0.1 mV, 10 ms. **(C-D)** Individual experiments (simple line) and group average (circles) fEPSPs normalized amplitude before (baseline, −10 to 0 min) and after (LTP, 30-40 min) TBS showing the lack of LTP in adult females (C; pubescent, P = 0.0017; adult, P = 0.4797). In contrast, LTP was induced at both pubescent and adult males BLA synapses (D; pubescent, P = 0.0003; adult, P = 0.0090). **(E-F)** Cumulative probability plot of percent LTP from individual experiments. Data are shown as mean ± SEM (A-D). Females: pubescent, n = 8; adult, n = 8. Males: pubescent, n = 9; adult, n=8. n = individual rats, one slice per rat. Data were analyzed via paired t test (C-D). *P<0.05. Color code: females, dark (pubescent) or light (adult) blue; males, dark (pubescent) or light (adult) orange.

In female BLA, while the TBS protocol effectively induced a lasting synaptic potentiation in slices obtained from pubescent rats, LTP was not induced by this protocol in slices prepared from adult rats (Fig. 10 A,C,E). In contrast, TBS-LTP was consistently expressed in both pubescent and adult male rats (Fig. 10 B,D), albeit to a lesser level at the adult stage compared to adolescent counterpart. Indeed, when plotting the cumulative distribution of LTP percent, data distribution of adults was left shifted compared to pubescent rats, therefore indicating a reduction of TBS-LTP at adulthood (Fig. 10F).

### Long-term depression appears at adulthood in the BLA of male rats

Long-term depression (LTD) is a widespread form of synaptic plasticity (Malenka & Bear, 2004) also present in the rat BLA. Indeed, low frequency stimulation (1hz for 15 min, LFS) at LA-BA synapses reliably induces LTD (S.-J. Wang & Gean, 1999). Thus, we systematically compared LFS-LTD in the BLA of pubescent and adult rats of both sexes (Fig. 11, Tab. 9). A 15-minute, 1Hz stimulation in BLA slices obtained from females in both age groups induces a robust LTD (Fig. 11 A,C,E). This protocol likewise elicited LTD in slices obtained from adult males (Fig. 11 B,D,F). To our surprise, LFS-LTD could not be induced in response to this protocol in pubescent males (Fig. 11 B,D,F). Taken together, the results suggest a degree of sex specificity to the maturational profile of LFS-LTD and TBS-LTP.

**Figure 11:**
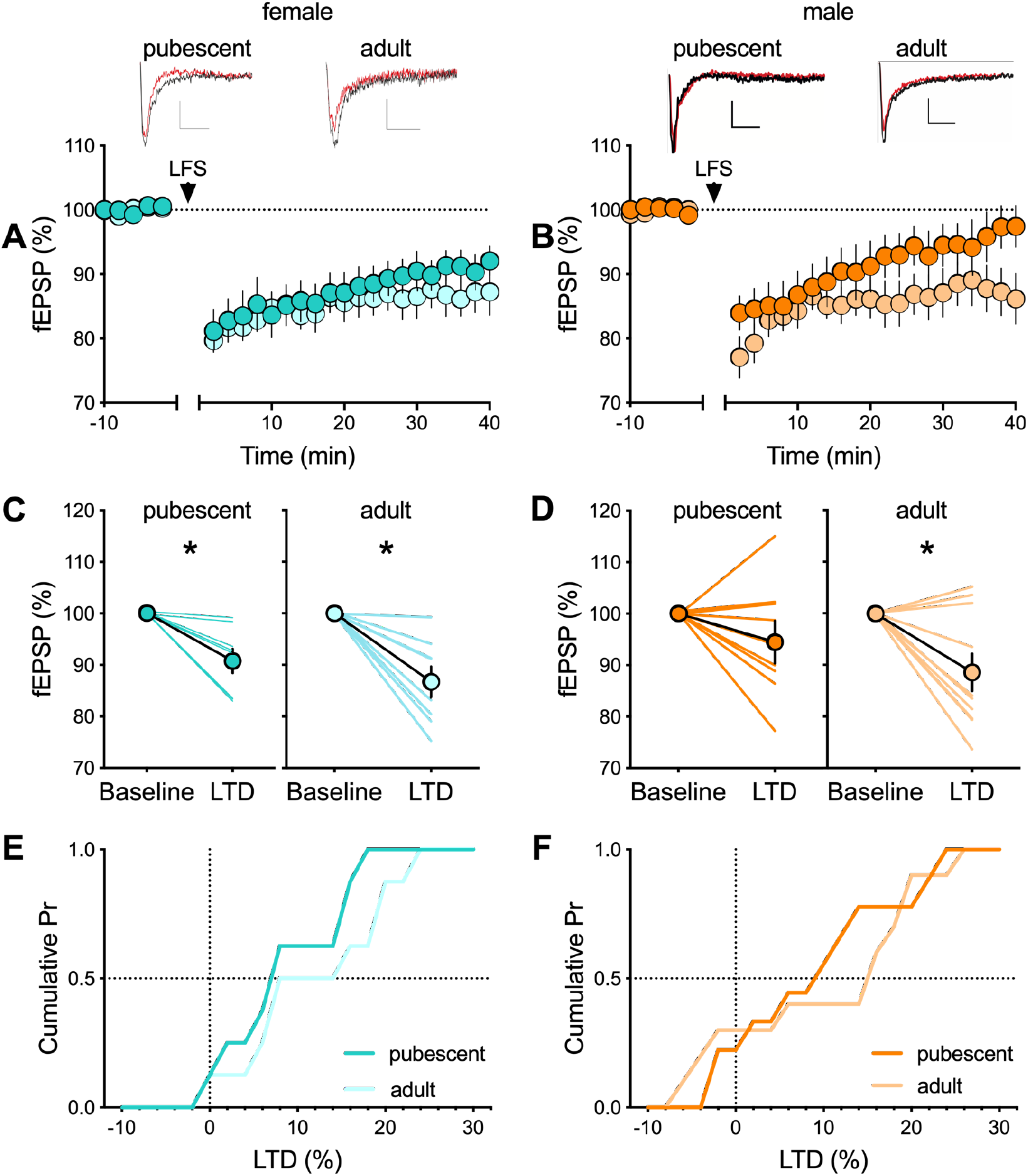
Long-term depression in the rat BLA. **(A-B)** Average time-courses of mean fEPSPs amplitude showing that low frequency stimulation, at 1Hz for 15 min (LFS, indicated arrow), induced LTD at BLA synapses in both pubescent and adult females (A). In contrast, LTD was present in BLA slices obtained from adult but not pubescent animals (B). Above: example traces, baseline (black) and 40 min post-stimulation (red). Scale bar: 0.1 mV, 10 ms. **(C-D)** Individual experiments (simple line) and group average (circles) fEPSPs’ normalized amplitude before (baseline, −10 to 0 min) and after (LTD, 30 to 40 min) LFS showing that LTD was present at both developmental stages in females (C; pubescent, P = 0.0139; adult, P = 0.0028). In contrast, LFS failed to induce a significant LTD in pubescent male slices, whereas it induced a significant LTD in adult male slices (D; pubescent, P = 0.2054; adult, P = 0.0118). **(E-F)** Cumulative probability plot of percent LTD from individual experiments. Data are shown as mean ± SEM (A-D). Females: pubescent, n = 7; adult, n = 8. Males: pubescent, n = 10; adult, n=9. n = individual rats, one slice per rat. Data were analyzed via paired t test (C-D). *P<0.05. Color code: females, dark (pubescent) or light (adult) blue; males, dark (pubescent) or light (adult) orange.

## Discussion

Cross-sectional portrayal of the morphology and neuronal properties of BLA neurons in rats of both sexes revealed sex-differences specific to adolescence and adulthood.

### Sex-differences in intrinsic properties of BLA neurons emerge at adulthood

We found that adult females had a higher rheobase, hyperpolarized membrane potential, and a lower propensity to trigger action potentials in response to depolarizing current steps than adult males. Consequently, principal neurons in the BLA of adult females are less excitable than those of adult males. Sex-differences in intrinsic properties of adult rats have already been described in other brain regions. Within the medial amygdala, a class of neurons exhibits lower spike frequency in response to depolarizing current in females than in males (Dalpian et al., 2019). In the medial nucleus of the central amygdala, Rouzer and Diaz also studied the sex-specific maturation of neurons intrinsic properties of adolescent and adult rats. While they found that males had lower AP thresholds than females suggesting more excitability in males, the quantification of spiking activity revealed that this was not directly translated in overall higher spiking activity in males than in females (Rouzer & Diaz, 2021). In the NAc core, medium spiny neurons’ intrinsic properties differ between males and females according to the estrous cycle phases (i.e. females are more excitable than males when in diestrus) (Proaño et al., 2018) and estradiol decrease the excitability of neurons from gonadectomized females (Proaño & Meitzen, 2020). Altogether, these results suggest that sex can profoundly influences principal neurons’ excitability in multiple brain areas. Blood hormone levels were not assessed directly, and while one cannot totally exclude that this may participate to experimental variability, the current observation that sex differences are not visible at puberty at a time when the estrous cycle of females is not well established supports the instrumental role of gonadal hormones.

Stress exposure induces sex-divergent effects in the amygdala: BA neurons’ firing rate augments in males and decreases in females (Blume et al., 2019). These changes have been attributed to stress-induced imbalance in excitation and inhibition (Blume et al., 2019; Mozhui et al., 2010; Rosenkranz et al., 2010; W. Zhang & Rosenkranz, 2012), and our results further suggest that initial sex-differences in neuronal intrinsic excitability could participate in the divergent effect of stress seen in males and females.

### Sex-differences in action potentials emerge at pubescence

Pubescent males had shorter action potentials with faster repolarization compared with females. An early study reported that sex does not influence the maturation of action potential waveform during the first post-natal month (Ehrlich et al., 2012). However, Guadagno and collaborators observed that juvenile (PND22-PND29) males have shorter action potentials with slower depolarization rates compared with females (Guadagno et al., 2020). Here, we found sex-differences reminiscent of those of pubescence (males still have shorter action potentials but faster repolarization). The current finding that the sex-differences disappear in adults suggests a profound sex-specific maturation of action potentials from prepuberty through late adolescence.

Pubescent females displayed smaller fAHP compared with males. Changes in the expression of large conductance KCa (BKCa) may explain these dissimilarities (Sah & Louise Faber, 2002). In the LA, specific blockade of BKCa leads to smaller fAHPs and slower repolarizations of action potentials. Furthermore, stress exposure recapitulates these effects and is associated with a down-regulation of BKCa channels (Guo et al., 2012). Hence, a down-regulation of BKCa channels in pubescent females compared with males could participate in sex-differences in fAHP but also in repolarization. The parallel finding that the overshoot also differ between pubescent males and females is coherent with an implication of these channels, the overshoot being determined by the opening of BKCa which are both Ca and voltage sensitive and initiate the repolarization phase.

Unlike fAHP, mAHP amplitude was not influenced by sex. However, age did influence the mAHP, with larger mAHPs in adult females compared with pubescents. The mAHP results from the activation of apamine-sensitive small conductance KCa channels (Sah & Louise Faber, 2002). Our results are therefore consistent with that idea of sex-specific profiles of SK channels expression during maturation.

Mixed evidence implicate the mAHP in the modulation of intrinsic spiking frequency in BLA neurons (Chen & Lang, 2003; Faber & Sah, 2002; Power & Sah, 2008). Although multiple factors could explain these discrepancies (the age of animals, in vivo vs in vitro, recording in LA or BA), smaller mAHP are consistently linked to an increased excitability. Furthermore, adolescent male rats that are responsive to restraint stress express higher spike frequency in both LA and BA neurons (Hetzel & Rosenkranz, 2014). While this increased excitability is accompanied by smaller mAHP in the LA, there was no effect on the mAHP in BA neurons. Thus, the mAHP is not constantly correlated with the spiking frequency in the BA. When we recorded BA neurons, the increased excitability seen in male adults compared females of the same age was not echoed by sex-differences in the mAHP. Nevertheless, in pubescent, high spiking frequency and smaller mAHP were concomitant (Table 1).

### Basic excitatory synaptic properties

We recently showed that the input-profile and the release probability of excitatory synapses to prefrontal cortex (PFC) pyramidal neurons were consistent between pubescence and adulthood in both sexes (Bernabeu et al., 2020). In the BLA, the input-output curves built in pubescent rats reached higher maxima than in adults, in support of the idea that the BLA reaches maturity late in life. Pre-synaptic (e.g. release probability) or post-synaptic (e.g. receptor trafficking, AMPAR and NMDAR expression; Hwang & Lupica, 2020) maturation of the LA-BA synapses could explain the dissimilarities between pubescent and adult rats. In the present study, the AMPA/NMDA ratio in females was stable through development, and thus this mechanism is unlikely to account for the maturation of the input-profile in female rats. The AMPA/NMDA ratio of pubescent males was higher compared with females of the same age and this sex-difference was not found in adults. Interestingly, the distribution of AMPA/NMDA ratios was bi-modal in pubescent but not adult males, perhaps an indication of a specific developmental signature in males. In contrast, previous work from our group conducted in the mice PFC demonstrated that AMPA/NMDA ratios remained stable between adolescence and adulthood (Iafrati et al., 2016). In the NAc (Kasanetz & Manzoni, 2009), AMPA/NMDA ratios are homogenous in adolescent mice while they express a bi-modal distribution in adults, contrary to what we observed in the BLA. Unfortunately, the studies did not include females. In any case, it is safe to conclude that the developmental trajectory of AMPA/NMDA ratios highly depends on the brain region.

The sex-specific maturation extended to sEPSCs whose amplitude augmented in adult females in keeping with the accompanying maturation of dendritic spines in this group (see below). sEPSCs’ properties remain overall comparable across sexes during maturation, suggesting that AMPA currents play a minor role in the high AMPA/NMDA ratios measured in pubescent males. In the BA of adult rats, mEPSCs’ frequency was increased when females were in diestrus but comparable to males while in proestrus (Blume et. al, 2017), leading to an overall increase in mEPSCs’ frequency in females compared with males, while mEPSCs’ amplitudes were similar. We did not observe such sex difference perhaps because of several methodological differences: our experiments were blind to the estrous cycle; we used coronal vs horizontal brain slices, we favored K^+^Gluconate internal medium over Cesium based solution, and we recorded spontaneous activity in the presence or absence of TTX.

Interestingly, Blume and coworkers correlated this sex-specific increase in frequency with an increase in spine density in females (independently of estrous cycle), while in our hands the lack of sex-differences in sEPSCs was accompanied by stable dendritic spine density. Thus, the studies are divergent with one another, but coherent on their own, pointing to the possibility that somewhat different populations of neurons/dendrites were recorded/reconstructed. In support of this idea, one must note that Blume and coworkers discarded non pyramidal shaped neurons while we exploited all glutamatergic principal neurons, from stellate to pyramidal (See methods).

### Dendritic spines

Our results replicate and extend upon previous morphological reconstructions of BLA principal neurons in male rats and mice showing that spine density remained stable between pubescence and adulthood (Bosch & Ehrlich, 2015; Ryan et al., 2016), by demonstrating that this remains the case in females. Although we elected not to formally classify spines into the classical categories (thin, stubby, mushroom), the majority of spines identified in our rat groups were clearly thin as previously showed in the BLA (J.-Y. Zhang et al., 2019). Quantitative analysis of our data revealed a remodeling of dendritic spines after puberty, particularly in females. Principal neurons of pubescent females had shorter spine lengths, but larger spine head diameters and spine volumes. In the BLA, large spine head diameters do not correlate with large EPSC amplitude (J.-Y. Zhang et al., 2019), in agreement with the current lack of difference in sEPSCs amplitude in adults of both sexes (see above).

### Sex influences the maturational profiles of LTP and LTD

The synaptic repertoire NMDAR is an important determinant of synaptic plasticity (Paoletti et al., 2013). Notably, in the BLA of young male rats (PND-17-24), GluN2A- and GluN2B-containing NMDAR subunits underlie LTP and LTD at auditory thalamic-LA pathway, respectively (Dalton et al., 2012). NMDAR activity or subunits mRNA expression can be modulated by gonadectomy in males (van den Buuse et al., 2017), estrous cycle in females (Picard et al., 2019), or sex (Damborsky & Winzer-Serhan, 2012; Qi et al., 2016). In the BLA, a recent study (Guadagno et al., 2020) did not report sex-differences in NMDAR and AMPAR subunit protein levels in pre-pubertal rats, but showed a sex-dependent shift of protein levels in response to stress. While the developmental decrease of synaptic GluN2B-containing NMDARs is completed by the second post-natal week in certain areas such as the hippocampus and the NAc (Bellone & Nicoll, 2007; Chavis & Westbrook, 2001; Kasanetz & Manzoni, 2009; Mierau et al., 2004), it is delayed towards adulthood in the PFC (Iafrati et al., 2016), suggesting that NMDAR composition can fluctuate late in development. Altogether, these data suggest that the sex-dependent divergent maturational trajectories of LTP and LTD could be influenced by sex-specific maturation of NMDAR subtypes in the BLA.

TBS specifically failed to induce LTP in the BLA of adult females. Compelling evidence indicate sex-differences in LTP and learning (Gall et al., 2021). Several studies reported stronger LTP in males in the hippocampus of adult mice, while a five-paired theta burst akin to ours failed to induce LTP in female though triggered a robust LTP in males; conversely, a 10-paired TBS protocol induced LTP in both sexes, suggesting a higher threshold for LTP induction in females (W. Wang et al., 2018). Thus, in the CNS, sex-differences in LTP are commonly observed and the BLA is no exception.

Finally, LFS-LTD was present throughout maturation in females, but only emerged at adulthood in males. Interestingly, GABAergic inhibition is necessary to induce LFS-LTD in the BLA (Rammes et al., 2001). Even though GABAergic basic synaptic properties are mature at adolescence (Ehrlich et al., 2013), there is evidence showing that BLA circuits are less inhibited in adolescent pre-pubertal male rats (-PND40) than in adults (Selleck et al., 2018; W. Zhang & Rosenkranz, 2012). In addition, early stress exposure enhances LFS-LTD in the BLA (Danielewicz & Hess, 2014). In the hippocampus, stress exposure only enhances LTD in males in early adolescence and this is concomitant with sex-differences in both basal level of circulating corticosteroids and glucocorticoid receptors expression (Huang et al., 2012). Thus, potential lines of research to elucidate the mechanisms preventing LTD in pubescent males include GABAergic system and glucocorticoids.

In conclusion, these data illustrate how male and female rats follow divergent maturational trajectories and identify previously unreported sex-differences at adulthood. Taken together with previous work (Blume et al., 2017, 2019; Hetzel & Rosenkranz, 2014), this study highlights the complexity of the cellular substrates to the BLA linked sex-specific behaviors at adolescence and adulthood.

Both a bigger range of behavioral studies as well as extended characterizations of plasticity and synaptic functions in animals of both sexes in other brain regions are necessary to better understand how sex-differences influence brain functions and behavior.

## Author contributions

Pauline Guily: Conceptualization; Data curation; Formal analysis; Validation; Writing: review and editing.

Olivier Lassalle: Data curation; Formal analysis; Validation; Methodology.

Pascale Chavis: Conceptualization; Methodology and editing.

Olivier JJ Manzoni: Conceptualization; Supervision; Funding acquisition; Methodology; Writing: original draft, review and editing; Project administration.

## Acknowledgements

The authors are grateful to the Chavis-Manzoni team members for helpful discussions and to Dr. A.F. Scheyer for critical reading and help with writing the manuscript.

## References

Bara, A., Manduca, A., Bernabeu, A., Borsoi, M., Serviado, M., Lassalle, O., Murphy, M., Wager-Miller, J., Mackie, K., Pelissier-Alicot, A.-L., Trezza, V., & Manzoni, O. J. (2018). Sex-dependent effects of in utero cannabinoid exposure on cortical function. ELife, 7. https://doi.org/10.7554/eLife.36234

Bellone, C., & Nicoll, R. A. (2007). Rapid Bidirectional Switching of Synaptic NMDA Receptors. Neuron, 55(5), 779–785. https://doi.org/10.1016/j.neuron.2007.07.035

Bender, R. A., Zhou, L., Vierk, R., Brandt, N., Keller, A., Gee, C. E., Schäfer, M. K. E., & Rune, G. M. (2017). Sex-Dependent Regulation of Aromatase-Mediated Synaptic Plasticity in the Basolateral Amygdala. The Journal of Neuroscience, 37(6), 1532–1545. https://doi.org/10.1523/JNEUROSCI.1532-16.2016

Bernabeu, A., Bara, A., Manduca, A., Borsoi, M., Lassalle, O., Pelissier-Alicot, A.-L., & Manzoni, O. J. (2020). Sex-specific maturational trajectory of endocannabinoid plasticity in the rat prefrontal cortex [Preprint]. Neuroscience. https://doi.org/10.1101/2020.10.09.332965

Blume, S. R., Freedberg, M., Vantrease, J. E., Chan, R., Padival, M., Record, M. J., DeJoseph, M. R., Urban, J. H., & Rosenkranz, J. A. (2017). Sex- and Estrus-Dependent Differences in Rat Basolateral Amygdala. The Journal of Neuroscience, 37(44), 10567–10586. https://doi.org/10.1523/JNEUROSCI.0758-17.2017

Blume, S. R., Padival, M., Urban, J. H., & Rosenkranz, J. A. (2019). Disruptive effects of repeated stress on basolateral amygdala neurons and fear behavior across the estrous cycle in rats. Scientific Reports, 9(1), 12292. https://doi.org/10.1038/s41598-019-48683-3

Bocchio, M., Nabavi, S., & Capogna, M. (2017). Synaptic Plasticity, Engrams, and Network Oscillations in Amygdala Circuits for Storage and Retrieval of Emotional Memories. Neuron, 94(4), 731–743. https://doi.org/10.1016/j.neuron.2017.03.022

Borsoi, M., Manduca, A., Bara, A., Lassalle, O., Pelissier-Alicot, A.-L., & Manzoni, O. J. (2019). Sex Differences in the Behavioral and Synaptic Consequences of a Single in vivo Exposure to the Synthetic Cannabimimetic WIN55,212-2 at Puberty and Adulthood. Frontiers in Behavioral Neuroscience, 13, 23. https://doi.org/10.3389/fnbeh.2019.00023

Bosch, D., & Ehrlich, I. (2015). Postnatal maturation of GABAergic modulation of sensory inputs onto lateral amygdala principal neurons: Development of sensory input inhibition in mouse lateral amygdala. The Journal of Physiology, 593(19), 4387–4409. https://doi.org/10.1113/JP270645

Bramen, J. E., Hranilovich, J. A., Dahl, R. E., Forbes, E. E., Chen, J., Toga, A. W., Dinov, I. D., Worthman, C. M., & Sowell, E. R. (2011). Puberty Influences Medial Temporal Lobe and Cortical Gray Matter Maturation Differently in Boys Than Girls Matched for Sexual Maturity. Cerebral Cortex, 21(3), 636–646. https://doi.org/10.1093/cercor/bhq137

Breslau, J., Gilman, S. E., Stein, B. D., Ruder, T., Gmelin, T., & Miller, E. (2017). Sex differences in recent first-onset depression in an epidemiological sample of adolescents. Translational Psychiatry, 7(5), e1139–e1139. https://doi.org/10.1038/tp.2017.105

Brinley-Reed, M., Mascagni, F., & McDonald, A. J. (1995). Synaptology of prefrontal cortical projections to the basolateral amygdala: An electron microscopic study in the rat. Neuroscience Letters, 202(1), 45–48. https://doi.org/10.1016/0304-3940(95)12212-5

Cahill, L., Uncapher, M., Kilpatrick, L., Alkire, M. T., & Turner, J. (2004). Sex-Related Hemispheric Lateralization of Amygdala Function in Emotionally Influenced Memory: An fMRI Investigation. Learning & Memory, 11(3), 261–266. https://doi.org/10.1101/lm.70504

Canli, T., Desmond, J. E., Zhao, Z., & Gabrieli, J. D. E. (2002). Sex differences in the neural basis of emotional memories. Proceedings of the National Academy of Sciences, 99(16), 10789–10794. https://doi.org/10.1073/pnas.162356599

Casey, B. J., Jones, R. M., & Hare, T. A. (2008). The Adolescent Brain. Annals of the New York Academy of Sciences, 1124, 111–126. https://doi.org/10.1196/annals.1440.010

Chaouloff, F., Hémar, A., & Manzoni, O. (2007). Acute Stress Facilitates Hippocampal CA1 Metabotropic Glutamate Receptor-Dependent Long-Term Depression. Journal of Neuroscience, 27(27), 7130–7135. https://doi.org/10.1523/JNEUROSCI.1150-07.2007

Chavis, P., & Westbrook, G. (2001). Integrins mediate functional pre- and postsynaptic maturation at a hippocampal synapse. Nature, 411(6835), 317–321. https://doi.org/10.1038/35077101

Chen, J. C. T., & Lang, E. J. (2003). Inhibitory control of rat lateral amygdaloid projection cells. Neuroscience, 121(1), 155–166. https://doi.org/10.1016/S0306-4522(03)00430-5

Christiansen, D. M., & Berke, E. T. (2020). Gender- and Sex-Based Contributors to Sex Differences in PTSD. Current Psychiatry Reports, 22(4), 19. https://doi.org/10.1007/s11920-020-1140-y

Dalpian, F., Rasia-Filho, A. A., & Calcagnotto, M. E. (2019). Sexual dimorphism, estrous cycle and laterality determine the intrinsic and synaptic properties of medial amygdala neurons in rat. Journal of Cell Science, 132(9), jcs227793. https://doi.org/10.1242/jcs.227793

Dalton, G. L., Wu, D. C., Wang, Y. T., Floresco, S. B., & Phillips, A. G. (2012). NMDA GluN2A and GluN2B receptors play separate roles in the induction of LTP and LTD in the amygdala and in the acquisition and extinction of conditioned fear. Neuropharmacology, 62(2), 797–806. https://doi.org/10.1016/j.neuropharm.2011.09.001

Damborsky, J. C., & Winzer-Serhan, U. H. (2012). Effects of sex and chronic neonatal nicotine treatment on NKCC1, KCC2, BDNF, NR2A and NR2B mRNA expression in the postnatal rat hippocampus. Neuroscience, 225, 105–117. https://doi.org/10.1016/j.neuroscience.2012.09.002

Danielewicz, J., & Hess, G. (2014). Early life stress alters synaptic modification range in the rat lateral amygdala. Behavioural Brain Research, 265, 32–37. https://doi.org/10.1016/j.bbr.2014.02.012

Daviu, N., Bruchas, M. R., Moghaddam, B., Sandi, C., & Beyeler, A. (2019). Neurobiological links between stress and anxiety. Neurobiology of Stress, 11, 100191. https://doi.org/10.1016/j.ynstr.2019.100191

Debanne, D., & Poo, M.-M. (2010). Spike-timing dependent plasticity beyond synapse—Pre- and post-synaptic plasticity of intrinsic neuronal excitability. Frontiers in Synaptic Neuroscience, 2, 21. https://doi.org/10.3389/fnsyn.2010.00021

Ehrlich, D. E., Ryan, S. J., Hazra, R., Guo, J.-D., & Rainnie, D. G. (2013). Postnatal maturation of GABAergic transmission in the rat basolateral amygdala. Journal of Neurophysiology, 110(4), 926–941. https://doi.org/10.1152/jn.01105.2012

Ehrlich, D. E., Ryan, S. J., & Rainnie, D. G. (2012). Postnatal development of electrophysiological properties of principal neurons in the rat basolateral amygdala: Development of principal neuron electrophysiology in the rat basolateral amygdala. The Journal of Physiology, 590(19), 4819–4838. https://doi.org/10.1113/jphysiol.2012.237453

Faber, E. S. L., Callister, R. J., & Sah, P. (2001). Morphological and Electrophysiological Properties of Principal Neurons in the Rat Lateral Amygdala In Vitro. Journal of Neurophysiology, 85(2), 714–723. https://doi.org/10.1152/jn.2001.85.2.714

Faber, E. S. L., & Sah, P. (2002). Physiological Role of Calcium-Activated Potassium Currents in the Rat Lateral Amygdala. Journal of Neuroscience, 22(5), 1618–1628. https://doi.org/10.1523/JNEUROSCI.22-05-01618.2002

Farb, C., Aoki, C., Milner, T., Kaneko, T., & LeDoux, J. (1992). Glutamate immunoreactive terminals in the lateral amygdaloid nucleus: A possible substrate for emotional memory. Brain Research, 593(2), 145–158. https://doi.org/10.1016/0006-8993(92)91303-V

Folkes, O. M., Báldi, R., Kondev, V., Marcus, D. J., Hartley, N. D., Turner, B. D., Ayers, J. K., Baechle, J. J., Misra, M. P., Altemus, M., Grueter, C. A., Grueter, B. A., & Patel, S. (2020). An endocannabinoid-regulated basolateral amygdala–nucleus accumbens circuit modulates sociability. The Journal of Clinical Investigation, 130(4), 1728–1742. https://doi.org/10.1172/JCI131752

Gall, C. M., Le, A. A., & Lynch, G. (2021). Sex differences in synaptic plasticity underlying learning. Journal of Neuroscience Research, 0(0), 1–19. https://doi.org/10.1002/jnr.24844

Geary, C. G., Wilk, V. C., Barton, K. L., Jefferson, P. O., Binder, T., Bhutani, V., Baker, C. L., Fernando-Peiris, A. J., Mousley, A. L., Rozental, S. F. A., Thompson, H. M., Touchon, J. C., Esteban, D. J., & Bergstrom, H. C. (2021). Sex differences in gut microbiota modulation of aversive conditioning, open field activity, and basolateral amygdala dendritic spine density. Journal of Neuroscience Research, 99(7), 1780–1801. https://doi.org/10.1002/jnr.24848

Goddings, A.-L., Mills, K. L., Clasen, L. S., Giedd, J. N., Viner, R. M., & Blakemore, S.-J. (2014). The influence of puberty on subcortical brain development. NeuroImage, 88, 242–251. https://doi.org/10.1016/j.neuroimage.2013.09.073

Greiner, E. M., Müller, I., Norris, M. R., Ng, K. H., & Sangha, S. (2019). Sex differences in fear regulation and reward-seeking behaviors in a fear-safety-reward discrimination task. Behavioural Brain Research, 368, 111903. https://doi.org/10.1016/j.bbr.2019.111903

Gruene, T. M., Flick, K., Stefano, A., Shea, S. D., & Shansky, R. M. (2015). Sexually divergent expression of active and passive conditioned fear responses in rats. eLife, 4, e11352. https://doi.org/10.7554/eLife.11352

Guadagno, A., Verlezza, S., Long, H., Wong, T. P., & Walker, C.-D. (2020). It Is All in the Right Amygdala: Increased Synaptic Plasticity and Perineuronal Nets in Male, But Not Female, Juvenile Rat Pups after Exposure to Early-Life Stress. The Journal of Neuroscience, 40(43), 8276–8291. https://doi.org/10.1523/JNEUROSCI.1029-20.2020

Guo, Y., Liu, S., Cui, G.-B., Ma, L., Feng, B., Xing, J., Yang, Q., Li, X., Wu, Y., Xiong, L., Zhang, W., & Zhao, M. (2012). Acute stress induces down-regulation of large-conductance Ca2+-activated potassium channels in the lateral amygdala. The Journal of Physiology, 590(4), 875–886. https://doi.org/10.1113/jphysiol.2011.223784

Hetzel, A., & Rosenkranz, J. A. (2014). Distinct Effects of Repeated Restraint Stress on Basolateral Amygdala Neuronal Membrane Properties in Resilient Adolescent and Adult Rats. Neuropsychopharmacology, 39(9), 2114–2130. https://doi.org/10.1038/npp.2014.60

Huang, C.-C., Chen, J.-P., Yeh, C.-M., & Hsu, K.-S. (2012). Sex difference in stress-induced enhancement of hippocampal CA1 long-term depression during puberty. Hippocampus, 22(7), 1622–1634. https://doi.org/10.1002/hipo.21003

Hunsberger, M. S., & Mynlieff, M. (2020). BK potassium currents contribute differently to action potential waveform and firing rate as rat hippocampal neurons mature in the first postnatal week. Journal of Neurophysiology, 124(3), 703–714. https://doi.org/10.1152/jn.00711.2019

Hwang, E.-K., & Lupica, C. R. (2020). Altered Corticolimbic Control of the Nucleus Accumbens by Long-term Δ9-Tetrahydrocannabinol Exposure. Biological Psychiatry, 87(7), 619–631. https://doi.org/10.1016/j.biopsych.2019.07.024

Iafrati, J., Malvache, A., Gonzalez Campo, C., Orejarena, M. J., Lassalle, O., Bouamrane, L., & Chavis, P. (2016). Multivariate synaptic and behavioral profiling reveals new developmental endophenotypes in the prefrontal cortex. Scientific Reports, 6, 35504. https://doi.org/10.1038/srep35504

Janak, P. H., & Tye, K. M. (2015). From circuits to behaviour in the amygdala. Nature, 517(7534), 284–292. https://doi.org/10.1038/nature14188

Kasanetz, F., & Manzoni, O. J. (2009). Maturation of Excitatory Synaptic Transmission of the Rat Nucleus Accumbens From Juvenile to Adult. Journal of Neurophysiology, 101(5), 2516–2527. https://doi.org/10.1152/jn.91039.2008

Kirshner, H., Aguet, F., Sage, D., & Unser, M. (2013). 3-D PSF fitting for fluorescence microscopy: Implementation and localization application. Journal of Microscopy, 249(1), 13–25. https://doi.org/10.1111/j.1365-2818.2012.03675.x

Krężel, W., Dupont, S., Krust, A., Chambon, P., & Chapman, P. F. (2001). Increased anxiety and synaptic plasticity in estrogen receptor β-deficient mice. Proceedings of the National Academy of Sciences, 98(21), 12278–12282. https://doi.org/10.1073/pnas.221451898

Kuehner, C. (2017). Why is depression more common among women than among men? The Lancet Psychiatry, 4(2), 146–158. https://doi.org/10.1016/S2215-0366(16)30263-2

Labouesse, M. A., Lassalle, O., Richetto, J., Iafrati, J., Weber-Stadlbauer, U., Notter, T., Gschwind, T., Pujadas, L., Soriano, E., Reichelt, A. C., Labouesse, C., Langhans, W., Chavis, P., & Meyer, U. (2017). Hypervulnerability of the adolescent prefrontal cortex to nutritional stress via reelin deficiency. Molecular Psychiatry, 22(7), 961–971. https://doi.org/10.1038/mp.2016.193

Lebron-Milad, K., & Milad, M. R. (2012). Sex differences, gonadal hormones and the fear extinction network: Implications for anxiety disorders. Biology of Mood & Anxiety Disorders, 2, 3. https://doi.org/10.1186/2045-5380-2-3

Mahan, A. L., & Ressler, K. J. (2012). Fear conditioning, synaptic plasticity and the amygdala: Implications for posttraumatic stress disorder. Trends in Neurosciences, 35(1), 24–35. https://doi.org/10.1016/j.tins.2011.06.007

Malenka, R. C., & Bear, M. F. (2004). LTP and LTD: An Embarrassment of Riches. Neuron, 44(1), 5–21. https://doi.org/10.1016/j.neuron.2004.09.012

Malinow, R., & Malenka, R. C. (2002). AMPA Receptor Trafficking and Synaptic Plasticity. Annual Review of Neuroscience, 25(1), 103–126. https://doi.org/10.1146/annurev.neuro.25.112701.142758

Manduca, A., Bara, A., Larrieu, T., Lassalle, O., Joffre, C., Layé, S., & Manzoni, O. J. (2017). Amplification of mGlu5-endocannabinoid signaling rescues behavioral and synaptic deficits in a mouse model of adolescent and adult dietary polyunsaturated fatty acid imbalance. Journal of Neuroscience, 37(29), 6851–6868. https://doi.org/10.1523/JNEUROSCI.3516-16.2017

Martin, H. G. S., Lassalle, O., Brown, J. T., & Manzoni, O. J. (2016). Age-Dependent Long-Term Potentiation Deficits in the Prefrontal Cortex of the Fmr1 Knockout Mouse Model of Fragile X Syndrome. Cerebral Cortex, 26(5), 2084–2092. https://doi.org/10.1093/cercor/bhv031

Martin, H. G. S., & Manzoni, O. (2014). Late onset deficits in synaptic plasticity in the valproic acid rat model of autism. Frontiers in Cellular Neuroscience, 0. https://doi.org/10.3389/fncel.2014.00023

Matos, H. Y., Hernandez-Pineda, D., Charpentier, C. M., Rusk, A., Corbin, J. G., & Jones, K. S. (2020). Sex Differences in Biophysical Signatures across Molecularly Defined Medial Amygdala Neuronal Subpopulations. ENeuro, 7(4). https://doi.org/10.1523/ENEURO.0035-20.2020

McLean, C. P., Asnaani, A., Litz, B. T., & Hofmann, S. G. (2011). Gender differences in anxiety disorders: Prevalence, course of illness, comorbidity and burden of illness. Journal of Psychiatric Research, 45(8), 1027–1035. https://doi.org/10.1016/j.jpsychires.2011.03.006

Mierau, S. B., Meredith, R. M., Upton, A. L., & Paulsen, O. (2004). Dissociation of experiencedependent and -independent changes in excitatory synaptic transmission during development of barrel cortex. Proceedings of the National Academy of Sciences, 101(43), 15518–15523. https://doi.org/10.1073/pnas.0402916101

Moyer, C. E., & Zuo, Y. (2018). Cortical dendritic spine development and plasticity: Insights from in vivo imaging. Current Opinion in Neurobiology, 53, 76–82. https://doi.org/10.1016/j.conb.2018.06.002

Mozhui, K., Karlsson, R.-M., Kash, T. L., Ihne, J., Norcross, M., Patel, S., Farrell, M. R., Hill, E. E., Graybeal, C., Martin, K. P., Camp, M., Fitzgerald, P. J., Ciobanu, D. C., Sprengel, R., Mishina, M., Wellman, C. L., Winder, D. G., Williams, R. W., & Holmes, A. (2010). Strain Differences in Stress Responsivity Are Associated with Divergent Amygdala Gene Expression and Glutamate-Mediated Neuronal Excitability. Journal of Neuroscience, 30(15), 5357–5367. https://doi.org/10.1523/JNEUROSCI.5017-09.2010

Nestler, E. J., Barrot, M., DiLeone, R. J., Eisch, A. J., Gold, S. J., & Monteggia, L. M. (2002). Neurobiology of Depression. Neuron, 34(1), 13–25. https://doi.org/10.1016/S0896-6273(02)00653-0

Paoletti, P., Bellone, C., & Zhou, Q. (2013). NMDA receptor subunit diversity: Impact on receptor properties, synaptic plasticity and disease. Nature Reviews Neuroscience, 14(6), 383–400. https://doi.org/10.1038/nrn3504

Pape, H.-C., & Pare, D. (2010). Plastic Synaptic Networks of the Amygdala for the Acquisition, Expression, and Extinction of Conditioned Fear. Physiological Reviews, 90(2), 419–463. https://doi.org/10.1152/physrev.00037.2009

Phelps, E. A. (2004). Human emotion and memory: Interactions of the amygdala and hippocampal complex. Current Opinion in Neurobiology, 14(2), 198–202. https://doi.org/10.1016/j.conb.2004.03.015

Picard, N., Takesian, A. E., Fagiolini, M., & Hensch, T. K. (2019). NMDA 2A receptors in parvalbumin cells mediate sex-specific rapid ketamine response on cortical activity. Molecular Psychiatry, 24(6), 828–838. https://doi.org/10.1038/s41380-018-0341-9

Power, J. M., & Sah, P. (2008). Competition between Calcium-Activated K+ Channels Determines Cholinergic Action on Firing Properties of Basolateral Amygdala Projection Neurons. Journal of Neuroscience, 28(12), 3209–3220. https://doi.org/10.1523/JNEUROSCI.4310-07.2008

Proaño, S. B., & Meitzen, J. (2020). Estradiol decreases medium spiny neuron excitability in female rat nucleus accumbens core. Journal of Neurophysiology, 123(6), 2465–2475. https://doi.org/10.1152/jn.00210.2020

Proaño, S. B., Morris, H. J., Kunz, L. M., Dorris, D. M., & Meitzen, J. (2018). Estrous cycle-induced sex differences in medium spiny neuron excitatory synaptic transmission and intrinsic excitability in adult rat nucleus accumbens core. Journal of Neurophysiology, 120(3), 1356–1373. https://doi.org/10.1152/jn.00263.2018

Przybysz, K. R., Gamble, M. E., & Diaz, M. R. (2021). Moderate adolescent chronic intermittent ethanol exposure sex-dependently disrupts synaptic transmission and kappa opioid receptor function in the basolateral amygdala of adult rats. Neuropharmacology, 188, 108512. https://doi.org/10.1016/j.neuropharm.2021.108512

Qi, X., Zhang, K., Xu, T., Yamaki, V. N., Wei, Z., Huang, M., Rose, G. M., & Cai, X. (2016). Sex Differences in Long-Term Potentiation at Temporoammonic-CA1 Synapses: Potential Implications for Memory Consolidation. PLOS ONE, 11(11), e0165891. https://doi.org/10.1371/journal.pone.0165891

Radley, J. J., Farb, C. R., He, Y., Janssen, W. G. M., Rodrigues, S. M., Johnson, L. R., Hof, P. R., LeDoux, J. E., & Morrison, J. H. (2007). Distribution of NMDA and AMPA receptor subunits at thalamo-amygdaloid dendritic spines. Brain Research, 1134, 87–94. https://doi.org/10.1016/j.brainres.2006.11.045

Rammes, G., Eder, M., Dodt, H.-U., Kochs, E., & Zieglgänsberger, W. (2001). Long-term depression in the basolateral amygdala of the mouse involves the activation of interneurons. Neuroscience, 107(1), 85–97. https://doi.org/10.1016/S0306-4522(01)00336-0

Rau, A. R., Chappell, A. M., Butler, T. R., Ariwodola, O. J., & Weiner, J. L. (2015). Increased Basolateral Amygdala Pyramidal Cell Excitability May Contribute to the Anxiogenic Phenotype Induced by Chronic Early-Life Stress. Journal of Neuroscience, 35(26), 9730–9740. https://doi.org/10.1523/JNEUROSCI.0384-15.2015

Rosenkranz, J. A., Venheim, E. R., & Padival, M. (2010). Chronic Stress Causes Amygdala Hyperexcitability in Rodents. Biological Psychiatry, 67(12), 1128–1136. https://doi.org/10.1016/j.biopsych.2010.02.008

Rouzer, S. K., & Diaz, M. R. (2021). Factors of sex and age dictate the regulation of GABAergic activity by corticotropin-releasing factor receptor 1 in the medial sub-nucleus of the central amygdala. Neuropharmacology, 189, 108530. https://doi.org/10.1016/j.neuropharm.2021.108530

Ryan, S. J., Ehrlich, D. E., & Rainnie, D. G. (2016). Morphology and dendritic maturation of developing principal neurons in the rat basolateral amygdala. Brain Structure and Function, 221(2), 839–854. https://doi.org/10.1007/s00429-014-0939-x

Sage, D., Donati, L., Soulez, F., Fortun, D., Schmit, G., Seitz, A., Guiet, R., Vonesch, C., & Unser, M. (2017). DeconvolutionLab2: An open-source software for deconvolution microscopy. Methods, 115, 28–41. https://doi.org/10.1016/j.ymeth.2016.12.015

Sah, P., & Louise Faber, E. S. (2002). Channels underlying neuronal calcium-activated potassium currents. Progress in Neurobiology, 66(5), 345–353. https://doi.org/10.1016/S0301-0082(02)00004-7

Scherf, K. S., Smyth, J. M., & Delgado, M. R. (2013). The amygdala: An agent of change in adolescent neural networks. Hormones and Behavior, 64(2), 298–313. https://doi.org/10.1016/j.yhbeh.2013.05.011

Scheyer, A. F., Borsoi, M., Pelissier-Alicot, A.-L., & Manzoni, O. J. J. (2020a). Maternal Exposure to the Cannabinoid Agonist WIN 55,12,2 during Lactation Induces Lasting Behavioral and Synaptic Alterations in the Rat Adult Offspring of Both Sexes. ENeuro, 7(5). https://doi.org/10.1523/ENEURO.0144-20.2020

Scheyer, A. F., Borsoi, M., Pelissier-Alicot, A.-L., & Manzoni, O. J. J. (2020b). Perinatal THC exposure via lactation induces lasting alterations to social behavior and prefrontal cortex function in rats at adulthood. Neuropsychopharmacology: Official Publication of the American College of Neuropsychopharmacology, 45(11), 1826–1833. https://doi.org/10.1038/s41386-020-0716-x

Scheyer, A. F., Borsoi, M., Wager-Miller, J., Pelissier-Alicot, A.-L., Murphy, M. N., Mackie, K., & Manzoni, O. J. J. (2019). Cannabinoid exposure via lactation in rats disrupts perinatal programming of the GABA trajectory and select early-life behaviors. Biological Psychiatry, Epub ahead of print. https://doi.org/10.1016/j.biopsych.2019.08.023

Schneider, M. (2013). Adolescence as a vulnerable period to alter rodent behavior. Cell and Tissue Research, 354(1), 99–106. https://doi.org/10.1007/s00441-013-1581-2

Selleck, R. A., Zhang, W., Samberg, H. D., Padival, M., & Rosenkranz, J. A. (2018). Limited prefrontal cortical regulation over the basolateral amygdala in adolescent rats. Scientific Reports, 8(1), 17171. https://doi.org/10.1038/s41598-018-35649-0

Shansky, R. M., & Murphy, A. Z. (2021). Considering sex as a biological variable will require a global shift in science culture. Nature Neuroscience, 24(4), 457–464. https://doi.org/10.1038/s41593-021-00806-8

Spear, L. P. (2000). The adolescent brain and age-related behavioral manifestations. Neuroscience & Biobehavioral Reviews, 24(4), 417–463. https://doi.org/10.1016/S0149-7634(00)00014-2

Thomazeau, A., Lassalle, O., Iafrati, J., Souchet, B., Guedj, F., Janel, N., Chavis, P., Delabar, J., & Manzoni, O. J. (2014). Prefrontal deficits in a murine model overexpressing the down syndrome candidate gene dyrk1a. The Journal of Neuroscience: The Official Journal of the Society for Neuroscience, 34(4), 1138–1147. https://doi.org/10.1523/JNEUROSCI.2852-13.2014

van den Buuse, M., Low, J. K., Kwek, P., Martin, S., & Gogos, A. (2017). Selective enhancement of NMDA receptor-mediated locomotor hyperactivity by male sex hormones in mice. Psychopharmacology, 234(18), 2727–2735. https://doi.org/10.1007/s00213-017-4668-8

Wang, S.-J., & Gean, P.-W. (1999). Long-Term Depression of Excitatory Synaptic Transmission in the Rat Amygdala. Journal of Neuroscience, 19(24), 10656–10663. https://doi.org/10.1523/JNEUROSCI.19-24-10656.1999

Wang, W., Le, A. A., Hou, B., Lauterborn, J. C., Cox, C. D., Levin, E. R., Lynch, G., & Gall, C. M. (2018). Memory-Related Synaptic Plasticity Is Sexually Dimorphic in Rodent Hippocampus. Journal of Neuroscience, 38(37), 7935–7951. https://doi.org/10.1523/JNEUROSCI.0801-18.2018

Washburn, M., & Moises, H. (1992). Electrophysiological and morphological properties of rat basolateral amygdaloid neurons in vitro. The Journal of Neuroscience, 12(10), 4066–4079. https://doi.org/10.1523/JNEUROSCI.12-10-04066.1992

Yang, R., Zhang, B., Chen, T., Zhang, S., & Chen, L. (2017). Postpartum estrogen withdrawal impairs GABAergic inhibition and LTD induction in basolateral amygdala complex via downregulation of GPR30. European Neuropsychopharmacology, 27(8), 759–772. https://doi.org/10.1016/j.euroneuro.2017.05.010

Zhang, J.-Y., Liu, T.-H., He, Y., Pan, H.-Q., Zhang, W.-H., Yin, X.-P., Tian, X.-L., Li, B.-M., Wang, X.-D., Holmes, A., Yuan, T.-F., & Pan, B.-X. (2019). Chronic Stress Remodels Synapses in an Amygdala Circuit-Specific Manner. Biological Psychiatry, 85(3), 189–201. https://doi.org/10.1016/j.biopsych.2018.06.019

Zhang, W., & Rosenkranz, J. A. (2012). Repeated restraint stress increases basolateral amygdala neuronal activity in an age-dependent manner. Neuroscience, 226, 459–474. https://doi.org/10.1016/j.neuroscience.2012.08.051

Zhang, Y., Garcia, E., Sack, A.-S., & Snutch, T. P. (2020). L-type calcium channel contributions to intrinsic excitability and synaptic activity during basolateral amygdala postnatal development. Journal of Neurophysiology, 123(3), 1216–1235. https://doi.org/10.1152/jn.00606.2019

