## Supplemental Figure for "Sex-specific divergent maturational trajectories in the postnatal rat basolateral amygdala"

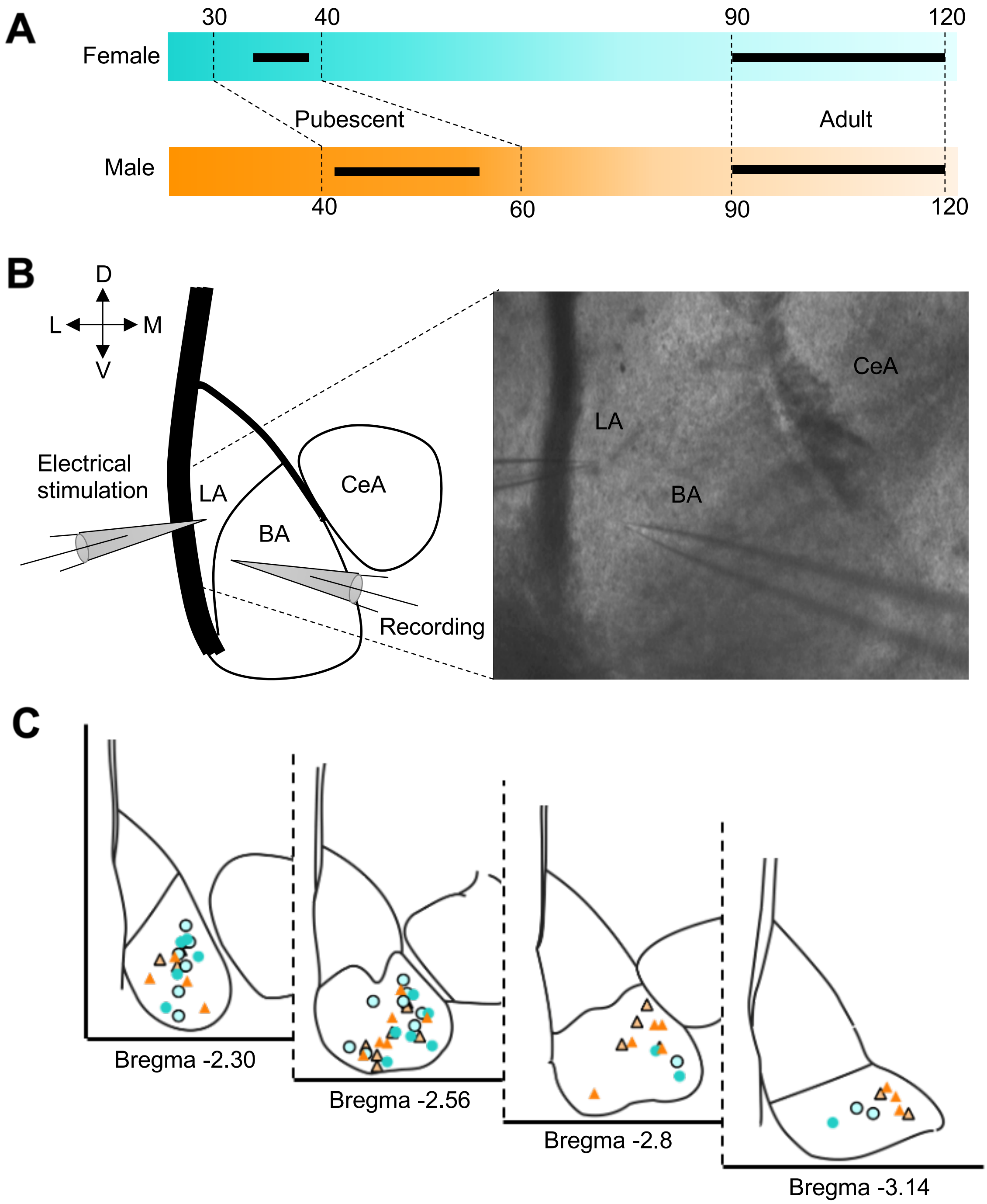
A
30
40
90
120
Female
Pubescent
Adult
Male
40
60
90
120
B
D
L
M
V
Electrical stimulation
LA
BA
CeA
Recording
CeA
LA
BA
C
Bregma -2.30
Bregma -2.56
Bregma -2.8
Bregma -3.14

**Supplementary figure 1: Age groups, location of electrophysiological recordings**

**(A)** Schematic representing the development of female (blue) and male (orange) rats from pubescence through adulthood. Female rats reach puberty earlier (30<PND<40) than males (40<PND<60) (Schneider, 2013). Black lines represent the age of rats used for our experiments (pubescent: female 33<P<38; male 41<P<56; adult 90<P<120 for both sexes). **(B)** Left, schematic representing the positions of the stimulating and recording electrodes during fEPSP recordings. The stimulating electrode was positioned in the lateral amygdala (LA), close to the external capsula (EC), dorsolateral to the recording electrode placed into the adjacent BA. Right, representative picture of electrodes' position during fEPSP recording. **(C)** Location of whole-cell patch-clamped BLA principal neurons during the recording of IV curves and sEPSCs.
