## Supplementary material for "Sex-specific divergent maturational trajectories in the postnatal rat basolateral amygdala": Tables

Values are mean ± SEM. F: Female, M: Male, Pub: Pubescent, Ad: Adult.

### Table 1.1 Intrinsic properties data – Two-way ANOVAs

| **Measure (unit)** | **Condition** | | **Value** | **Two-way ANOVA** |
|---|---|---|---|---|
| **RMP (mV)** | **F** | Pub | -67.96 ± 0.4775 N=17 | F (sex x dev 1, 75) = 0.3157 P = 0.5759 |
| | | Ad | -67.76 ± 0.4888 N=21 | **F (sex 1, 75) = 8.224 P = 0.0054** |
| | **M** | Pub | -66.68 ± 0.6145 N=22 | F (dev 1, 75) = 0.8427 P = 0.3616 |
| | | Ad | -65.86 ± 0.5777 N=19 | |
| **Rheobase (pA)** | **F** | Pub | 164.7 ± 9.120 N=17 | **F (sex x dev 1, 74) = 4.154 P=0.0451** |
| | | Ad | 174.3 ± 7.825 N=21 | F (sex 1, 74) = 0.7296 P=0.3958 |
| | **M** | Pub | 178.1 ± 13.66 N=21 | F (dev 1, 74) = 1.419 P=0.2374 |
| | | Ad | 141.6 ± 12.85 N=19 | |
| **Total number of spikes** | **F** | Pub | 180.059 ± 5.717 N=17 | **F (sex x dev 1, 73) = 9.085 P=0.0035** |
| | | Ad | 149.714 ± 7.070 N=21 | F (sex 1, 73) = 1.368 P=0.2460 |
| | **M** | Pub | 164.476 ± 8.890 N=21 | F (dev 1, 73) = 0.3341 P=0.5650 |
| | | Ad | 185.055 ± 10.895 N=18 | |

### Table 1.1 Intrinsic properties data - Post-hoc tests

| | **Sidak's multiple comparison test** | | | |
|---|---|---|---|---|
| **Measure (unit)** | **F: Pub vs Ad** | **M: Pub vs Ad** | **Pub: M vs F** | **Ad: M vs F** |
| **RMP (mV)** | P = 0.9622 | P = 0.4941 | P = 0.2089 | **P = 0.0330** |
| **Rheobase (pA)** | P = 0.8036 | **P = 0.0458** | P = 0.6543 | P = 0.0809 |
| **Total # spikes** | **P = 0.0276** | P = 0.1660 | P = 0.3594 | **P = 0.0078** |

### Table 2.1 Action potential properties - Two-way ANOVA

| Test – Measure (unit) | Condition | | Value | Two-way ANOVA |
|---|---|---|---|---|
| **AP threshold (mV)** | **F** | Pub | -39.69 ± 0.7882 N=14 | F (sex x dev 1, 59) = 0.7062 P=0.404<br>F (sex 1, 59) = 2.087 P=0.1539<br>F (dev 1, 59) = 0.06688 P=0.7968 |
| | | Ad | -39.27 ± 0.8737 N=16 | |
| | **M** | Pub | -40.13 ± 0.4649 N=16 | |
| | | Ad | -40.92 ± 0.6905 N=17 | |
| **AP amplitude (mV)** | **F** | Pub | 88.53 ± 1.907 N=14 | F (sex x dev 1, 59) = 3.015 P=0.0877<br>F (sex 1, 59) = 1.307 P=0.2576<br>F (dev 1, 59) = 0.1274 P=0.7224 |
| | | Ad | 85.91 ± 2.103 N=16 | |
| | **M** | Pub | 83.06 ± 1.557 N=16 | |
| | | Ad | 87.04 ± 1.954 N=17 | |
| **AP overshoot (mV)** | **F** | Pub | 48.79 ± 1.455 N=14 | F (sex x dev 1, 59) = 2.495 P=0.1195<br>F (sex 1, 59) = 3.573 P=0.0637<br>F (dev 1, 59) = 0.1067 P=0.7451 |
| | | Ad | 46.63 ± 1.671 N=16 | |
| | **M** | Pub | 42.80 ± 1.640 N=16 | |
| | | Ad | 46.09 ± 1.967 N=17 | |
| **AP duration (ms)** | **F** | Pub | 2.381 ± 0.0932 N=14 | F (sex x dev 1, 59) = 3.016 P=0.0876<br>F (sex 1, 59) = 2.166 P=0.1464<br>F (dev 1, 59) = 0.1788 P=0.6739 |
| | | Ad | 2.161 ± 0.0950 N=16 | |
| | **M** | Pub | 2.054 ± 0.0969 N=16 | |
| | | Ad | 2.188 ± 0.1151 N=17 | |
| **Half-width (ms)** | **F** | Pub | 1.046 ± 0.03523 N=14 | F (sex x dev 1, 59) = 3.015 P=0.0877<br>F (sex 1, 59) = 1.307 P=0.2576<br>F (dev 1, 59) = 0.1274 P=0.7224 |
| | | Ad | 0.9506 ± 0.04366 N=16 | |
| | **M** | Pub | 0.9281 ± 0.04343 N=16 | |
| | | Ad | 0.9406 ± 0.05602 N=17 | |
| **Depolarisation time (ms)** | **F** | Pub | 0.7650 ± 0.0255 N=14 | F (sex x dev 1, 59) = 0.1180 P=0.7325<br>F (sex 1, 59) = 0.001572 P=0.9685<br>F (dev 1, 59) = 0.9410 P=0.3360 |
| | | Ad | 0.7856 ± 0.0329 N=16 | |
| | **M** | Pub | 0.7550 ± 0.0270 N=16 | |
| | | Ad | 0.7982 ± 0.0406 N=17 | |
| **Repolarisation time (ms)** | **F** | Pub | 1.616 ± 0.0772 N=14 | **F (sex x dev 1, 56) = 6.918 P=0.0110**<br>**F (sex 1, 56) = 5.034 P=0.0288**<br>F (dev 1, 56) = 1.986 P=0.1643 |
| | | Ad | 1.311 ± 0.0544 N=15 | |
| | **M** | Pub | 1.299 ± 0.0840 N=16 | |
| | | Ad | 1.325 ± 0.0628 N=17 | |
| **Rise time (ms)** | **F** | Pub | 0.2574 ± 0.01114 N=14 | F (sex x dev 1, 59) = 0.389 P=0.5352<br>F (sex 1, 59) = 0.2930 P=0.5903<br>F (dev 1, 59) = 1.275 P=0.2635 |
| | | Ad | 0.2349 ± 0.01320 N=16 | |
| | **M** | Pub | 0.2424 ± 0.01230 N=16 | |
| | | Ad | 0.2359 ± 0.01387 N=17 | |
| **Decay time (ms)** | **F** | Pub | 0.4764 ± 0.01627 N=14 | F (sex x dev 1, 59) = 3.146 P=0.0813<br>F (sex 1, 59) = 2.939 P=0.0917<br>F (dev 1, 59) = 0.8244 P=0.3676 |
| | | Ad | 0.4207 ± 0.01898 N=16 | |
| | **M** | Pub | 0.4039 ± 0.02374 N=16 | |
| | | Ad | 0.4219 ± 0.02184 N=17 | |

**Table 2.2. Action potential properties – post-hoc test**

| | **Sidak's multiple comparison test** | | | |
|---|---|---|---|---|
| **Measure (unit)** | **F: Pub vs Ad** | **M: Pub vs Ad** | **Pub: M vs F** | **Ad: M vs F** |
| **AP threshold (mV)** | P = | P = | P = | **P =** |
| **AP amplitude (mV)** | P = 0.5709 | P = 0.2506 | P = 0.1004 | P = 0.8902 |
| **AP overshoot (mV)** | P = 0.6286 | P = 0.3145 | **P = 0.0392** | P = 0.9685 |
| **AP duration (ms)** | P = 0.2627 | P = 0.5704 | P = 0.0604 | P = 0.9769 |
| **Half-width (ms)** | P = 0.1601 | P = 0.9846 | P = 0.9727 | P = 0.9846 |
| **Depol duration (ms)** | P = 0.8891 | P = 0.5706 | P = 0.9727 | P = 0.9525 |
| **Repol duration (ms)** | **P = 0.0152** | P = 0.6103 | **P = 0.0025** | P = 0.9527 |
| **Rise time (ms)** | P = 0.4089 | P = 0.9190 | P = 0.6688 | P = 0.9977 |
| **Decay time (ms)** | P = 0.1335 | P = 0.7814 | **P = 0.0380** | P = 0.9988 |

**Table 3.1. AHP properties – two-way ANOVA**

| **Test – Measure (unit)** | **Condition** | | **Value** | |
|---|---|---|---|---|
| **fAHP amplitude (mV)** | **F** | Pub | $-4.473 \pm 0.5803$ N=12 | F (sex x dev 1, 54) = 3.720 P=0.0590 |
| | | Ad | $-5.793 \pm 0.5210$ N=15 | **F (sex 1, 54) = 5.708 P=0.0204** |
| | **M** | Pub | $-7.747 \pm 1.034$ N=14 | F (dev 1, 54) = 0.03558 P=0.8511 |
| | | Ad | $-6.142 \pm 0.7349$ N=17 | |
| **Time to reach fAHP (ms)** | **F** | Pub | $0.8158 \pm 0.05779$ N=12 | F (sex x dev 1, 54) = 1.199 P=0.2783 |
| | | Ad | $0.8450 \pm 0.09886$ N=14 | F (sex 1, 54) = 0.4541 P=0.5033 |
| | **M** | Pub | $0.8644 \pm 0.05324$ N=15 | F (dev 1, 54) = 0.1553 P=0.6951 |
| | | Ad | $0.7419 \pm 0.05782$ N=17 | |
| **mAHP amplitude (mV)** | **F** | Pub | $-6.797 \pm 0.8187$ N=12 | F (sex x dev 1, 56) = 2.763 P=0.1020 |
| | | Ad | $-10.35 \pm 0.7579$ N=16 | F (sex 1, 56) = 0.2171 P=0.6431 |
| | **M** | Pub | $-7.769 \pm 0.7879$ N=15 | **F (dev 1, 56) = 7.364 P=0.0088** |
| | | Ad | $-8.624 \pm 0.8335$ N=17 | |
| **Time to reach mAHP (ms)** | **F** | Pub | $62.29 \pm 5.277$ N=12 | F (sex x dev 1, 56) = 0.0299 P=0.863 |
| | | Ad | $69.65 \pm 4.681$ N=16 | F (sex 1, 56) = 1.933 P=0.1700 |
| | **M** | Pub | $53.29 \pm 5.994$ N=15 | F (dev 1, 56) = 2.108 P=0.1521 |
| | | Ad | $62.64 \pm 6.326$ N=17 | |

### Table 3.2 AHP properties - Post-hoc tests

| | Sidak's multiple comparison test | | | |
|---|---|---|---|---|
| **Measure (unit)** | **F: Pub vs Ad** | **M: Pub vs Ad** | **Pub: M vs F** | **Ad: M vs F** |
| **fAHP amplitude (mV)** | P = 0.4217 | P = 0.2366 | **P = 0.0106** | P = 0.9283 |
| **Time to reach fAHP (ms)** | P = 0.8626 | P = 0.4911 | P = 0.9508 | P = 0.3461 |
| **mAHP amplitude (mV)** | **P = 0.0084** | P = 0.6895 | P = 0.6688 | P = 0.2206 |
| **Time to reach mAHP (ms)** | P = 0.6242 | P = 0.4182 | P = 0.5067 | P = 0.5990 |

### Table 4.1 fEPSP input-output data - Two-way ANOVA

| **Measure (unit)** | **Condition** | | **Value** | **Two-way ANOVA** |
|---|---|---|---|---|
| **Maximum fEPSP amplitude (mV)** | **F** | Pub | 0.4715 ± 0.0229 N=10 | F (sex x dev 1, 32) = 1.666 P = 0.2060 |
| | | Ad | 0.3911 ± 0.0176 N=8 | F (sex 1, 32) = 3.521 P = 0.0697 |
| | **M** | Pub | 0.5593 ± 0.0338 N=8 | **F (dev 1, 32) = 17.59 P = 0.0002** |
| | | Ad | 0.4073 ± 0.0316N=10 | |

### Table 4.2 fEPSP input-output data - Post hoc tests

| | Sidak's multiple comparison test | | | |
|---|---|---|---|---|
| **Measure (unit)** | **F: Pub vs Ad** | **M: Pub vs Ad** | **Pub: M vs F** | **Ad: M vs F** |
| **Maximum fEPSP amplitude (mV)** | P = 0.0943 | **P = 0.0010** | P = 0.0634 | P = 0.8986 |

### Table 5.1. AMPA/NMDA ratio data – two-way ANOVA

| **Test – Measure (unit)** | **Condition** | | **Value** | **Two-way ANOVA** |
|---|---|---|---|---|
| **AMPA/NMDA ratio** | **F** | Pub | 0.6181 ± 0.06533 N=12 | **F (sex x dev 1, 40) = 4.739 P=0.0355** |
| | | Ad | 0.7712 ± 0.07989 N=9 | **F (sex 1, 40) = 4.769 P=0.0349** |
| | **M** | Pub | 1.049 ± 0.1373 N=12 | F (dev 1, 40) = 0.3929 P=0.5343 |
| | | Ad | 0.7719 ± 0.08110 N=11 | |

### Table 5.2 AMPA/NMDA ratio data - Post-hoc tests

| | Sidak's multiple comparison test | | | |
|---|---|---|---|---|
| **Measure (unit)** | **F: Pub vs Ad** | **M: Pub vs Ad** | **Pub: M vs F** | **Ad: M vs F** |
| **AMPA/NMDA ratio** | P = 0.4992 | P = 0.0939 | **P = 0.0048** | P > 0.9999 |

### Table 6.1 sEPSC properties data – two-way ANOVA

| Measure (unit) | Condition | | Value | Two-way ANOVA |
|---|---|---|---|---|
| **sEPSC frequency (Hz)** | **M** | Pub | 4.514 ± 0.3845 N=10 | F (sex x dev 1, 34) = 0.5828 P=0.4505<br>F (sex 1, 34) = 1.483e-005 P=0.9969<br>F (dev 1, 34) = 0.1607 P=0.6910 |
| | | Ad | 5.001 ± 0.4688 N=9 | |
| | **F** | Pub | 4.835 ± 0.3826 N=10 | |
| | | Ad | 4.683 ± 0.4421 N=9 | |
| **sEPSC amplitude (pA)** | **M** | Pub | 17. 46 ± 1.381 N=10 | F (sex x dev 1, 34) = 1.434 P=0.2394<br>F (sex 1, 34) = 0.4593 P=0.5025<br>F (dev 1, 34) = 3.076 P=0.0885 |
| | | Ad | 18.09 ± 1.511 N=9 | |
| | **F** | Pub | 15.32 ± 0.7851 N=10 | |
| | | Ad | 18.68 ± 0.5933 N=9 | |
| | | F | N=9 | |

### Table 6.2 sEPSC properties data – post-hoc tests

| | **Sidak's multiple comparison test** | | | |
|---|---|---|---|---|
| **Measure (unit)** | **F: Pub vs Ad** | **M: Pub vs Ad** | **Pub: M vs F** | **Ad: M vs F** |
| **sEPSC frequency (Hz)** | P = 0.9597 | P = 0.6590 | P = 0.8244 | P = 0.8432 |
| **sEPSC amplitude (pA)** | P = 0.0870 | P = 0.9079 | P = 0.3310 | P = 0.9230 |

### Table 7.1 Dendritic spine density – two-way ANOVA

| Measure (unit) | Condition | | Value | Two-way ANOVA |
|---|---|---|---|---|
| **Spine density (1/10μm)** | **M** | Pub | 17.77 ± N=8 | F (sex x dev 1, 26) = 0.4965 P=0.4873<br>F (sex 1, 26) = 0.8264 P=0.3717<br>F (dev 1, 26) = 0.4236 P=0.5209 |
| | | Ad | 17.67 ± 1.545 N=8 | |
| | **F** | Pub | 14.83 ± 1.476 N=7 | |
| | | Ad | 17.30 ± 1.236 N=7 | |

### Table 7.2 Dendritic spine density – post-hoc tests

| | **Sidak's multiple comparison test** | | | |
|---|---|---|---|---|
| **Measure (unit)** | **F: Pub vs Ad** | **M: Pub vs Ad** | **Pub: M vs F** | **Ad: M vs F** |
| **Spine density (1/10μm)** | P = 0.5929 | P = 0.9990 | P = 0.4587 | P = 0.9870 |

### Table 7.3 Dendritic spine morphology data – Kruskal-Wallis tests

| Measure (unit) | Condition | | Value | Kruskal-Wallis test |
|---|---|---|---|---|
| **Spine length (μm)** | **M** | Pub | 1.257 ± 0.01079 N=4658 | **P < 0.0001** |
| | | Ad | 1.206 ± 0.01067 N=4639 | |
| | **F** | Pub | 1.261 ± 0.01514 N=2450 | |
| | | Ad | 1.365 ± 0.01450 N=3908 | |
| **Spine head diameter (μm)** | **M** | Pub | 0.3264 ± 0.001784 N=4640 | **P < 0.0001** |
| | | Ad | 0.3284 ± 0.001928 N=4585 | |
| | **F** | Pub | 0.3472 ± 0.002399 N=2450 | |
| | | Ad | 0.3213 ± 0.001678 N=3784 | |
| **Spine volume ($\mu m^3$)** | **M** | Pub | 0.08388 ± 0.001039 N=4586 | **P = 0.0002** |
| | | Ad | 0.08596 ± 0.001131 N=4540 | |
| | **F** | Pub | 0.08953 ± 0.001536 N=2430 | |
| | | Ad | 0.08053 ± 0.001105 N=3769 | |

### Table 7.4 Dendritic spine morphology data – post-hoc tests

| | **Sidak's multiple comparison test** | | | |
|---|---|---|---|---|
| **Measure (unit)** | **F: Pub vs Ad** | **M: Pub vs Ad** | **Pub: M vs F** | **Ad: M vs F** |
| **Spine length (μm)** | **P = 0.0061** | **P = 0.0018** | P > 0.9999 | **P < 0.0001** |
| **Spine head diameter (μm)** | **P < 0.0001** | P > 0.9999 | **P < 0.0001** | P > 0.9999 |
| **Spine volume ($\mu m^3$)** | **P = 0.0002** | P > 0.9999 | **P = 0.0018** | P > 0.9999 |

### Table 8. TBS-LTP data

| Measure (unit) | Condition | | Value | Paired t test |
|---|---|---|---|---|
| **TBS-LTP**<br>Normalized fEPSP<br>0-10 min baseline (%) vs<br>30-40min post-tetanus (%) | **F** | Pub | 114.5 ± 2.915 N = 8 | **P = 0.0017** |
| | | Ad | 102.4 ± 3.173 N = 8 | P = 0.4797 |
| | **M** | Pub | 117.0 ± 2.853 N = 9 | **P = 0.0003** |
| | | Ad | 109.2 ± 2.565 N = 8 | **P = 0.0090** |

### Table 9. LFS-LTD data

| Measure (unit) | Condition | | Value | Paired t test |
|---|---|---|---|---|
| **LFS-LTD**<br>Normalized fEPSP<br>0-10 min baseline (%) vs<br>30-40min post-tetanus (%) | **F** | Pub | 91.82 ± 2.432 N = 7 | **P = 0.0139** |
| | | Ad | 86.71 ± 2.968 N = 8 | **P = 0.0028** |
| | **M** | Pub | 94.95 ± 3.676 N = 9 | P = 0.2054 |
| | | Ad | 88.58 ± 3.635 N = 10 | **P = 0.0118** |
